# The 3’ exonuclease TOE1 selectively processes snRNAs through recognition of Sm complex assembly and 5’ cap trimethylation

**DOI:** 10.1101/2023.08.15.553431

**Authors:** Tiantai Ma, Erica S Xiong, Rea M Lardelli, Jens Lykke-Andersen

## Abstract

Competing exonucleases that promote 3’ end maturation or degradation direct quality control of small non-coding RNAs, but how these enzymes distinguish normal from aberrant RNAs is poorly understood. The Pontocerebellar Hypoplasia 7 (PCH7)-associated 3’ exonuclease TOE1 promotes maturation of canonical small nuclear RNAs (snRNAs). Here, we demonstrate that TOE1 achieves specificity towards canonical snRNAs by recognizing Sm complex assembly and cap trimethylation, two features that distinguish snRNAs undergoing correct biogenesis from other small non-coding RNAs. Indeed, disruption of Sm complex assembly via snRNA mutations or protein depletions obstructs snRNA processing by TOE1, and in vitro snRNA processing by TOE1 is stimulated by a trimethylated cap. An unstable snRNA variant that normally fails to undergo maturation becomes fully processed by TOE1 when its degenerate Sm binding motif is converted into a canonical one. Our findings uncover the molecular basis for how TOE1 distinguishes snRNAs from other small non-coding RNAs and explain how TOE1 promotes maturation specifically of canonical snRNAs undergoing proper processing.

## Introduction

Small non-coding (nc)RNAs play key roles at all levels of gene regulation^1–7^. In eukaryotes, the majority of small ncRNAs are transcribed as precursor RNAs that need to undergo further processing to become functional mature RNAs, including trimming of 3’ end extensions^8,9^. These extensions serve as central hubs for quality control, where competing 3’-to-5’ exonucleases that promote maturation or degradation dictate whether the small ncRNAs undergo processing to functional molecules^10–13^ or are subjected to decay^12,14,15^. The competition between these processes can be influenced by post-transcriptional oligo(A) or –(U) tailing by terminal nucleotidyl transferases^12,16–19^. A key question is what dictates the specificity of the exonucleases and terminal nucleotidyl transferases that act on small ncRNAs to ultimately control their fate. A prevailing hypothesis is that enzymes that promote maturation recognize specific features of RNAs undergoing proper biogenesis, while more promiscuous decay enzymes degrade those RNAs that fail to conform to canonical processing. This predicts that maturation enzymes recognize common features of canonical RNAs that signify their correct biogenesis.

Recent studies have uncovered several 3’ to 5’ exonucleases that promote deadenylation and 3’ end maturation of human small ncRNAs. Two such exonucleases are the DEDD family deadenylases target of EGR1 protein 1 (TOE1; also known as CAF1Z)^20^ and poly(A)-specific ribonuclease (PARN)^21^. While TOE1 and PARN are homologous proteins and are both activated by oligo(A) tails^20,21^, they differ in their specificity for small ncRNAs. PARN has been reported to process 3’ ends of several types of small ncRNAs, including small nucleolar RNAs (snoRNAs)^11,22^ and the telomerase RNA component (TERC)^23–25^. By contrast, TOE1 is known to process 3’ ends of RNA Polymerase II-transcribed small nuclear RNAs (Pol II snRNAs; i.e. all snRNAs except U6 and U6atac)^10,12^, and has also been reported to process some snoRNAs, scaRNAs, and TERC^11,26^. Yet another DEDD deadenylase, the germ-cell specific PNLDC1, has been implicated in processing of Piwi-interacting (pi)RNAs^27–29^, and USB1, an exonuclease of the 2H phosphodiesterase family, is known to process U6 and U6atac snRNAs and miRNAs^30–32^. Functions of these enzymes are central to human health, with TOE1 mutations associated with *Pontocerebellar Hypoplasia Type 7* (PCH7)^10^, PARN mutations with *Dyskeratosis Congenita* and other human disorders^33,34^, PNLDC1 mutations with *azoospermia*^35^, and USB1 mutations with *Poikiloderma with neutropenia*^36^. Despite their importance in RNA metabolism and human disease, little is known about the molecular basis for the RNA specificity of these deadenylases.

SnRNAs are central components of the spliceosome that carries out pre-mRNA splicing. The maturation of Pol II snRNAs includes multiple steps in the nucleus and cytoplasm^37^. During transcription, a 7-methyl guanosine (m7G) cap is co-transcriptionally added to the snRNA 5’ end while the 3’ end is cleaved by the Integrator complex, leaving a short 3’ end tail^38,39^. The m7G cap subsequently promotes nuclear export via the export factor PHAX in association with the nuclear cap-binding complex (CBC)^40^. In the cytoplasm, the Sm complex is loaded onto the snRNAs with help of the SMN-Gemin2 complex, and protein arginine methyl transferases (PRMTs)^41–49^, and the m7G cap is hypermethylated to a trimethylguanosine (TMG) cap by trimethylguanosine synthase 1 (TGS1)^50–52^. The Sm complex and the TMG cap are subsequently recognized by Snurportin (SNUPN), which promotes nuclear import via Importin β^53,54^. Following nuclear import, the snRNAs undergo nucleotide modifications directed by scaRNAs and assemble with other snRNA-protein complexes (snRNPs) to form the spliceosome^55^.

Evidence suggests that TOE1 acts on Pol II snRNAs at least twice during their maturation^12^. During early biogenesis, prior to or upon nuclear export, TOE1 partially trims Pol II snRNA 3’ ends in a process that involves oligoadenylation^12^. The remainder of the tail is removed later, during or after nuclear import, when TOE1 acts on snRNAs a second time^12,56^. While TOE1 acts on all canonical Pol II snRNAs, tested unstable snRNA variants transcribed from snRNA pseudogenes and mutant U1 snRNA deleted of its Sm binding site escape 3’ end processing and are instead subjected to decay by degrading exonucleases^12,19,56,57^. The features of canonical snRNAs recognized by TOE1 during early and late processing steps and how TOE1 distinguishes canonical snRNAs from unstable snRNA variants and other small ncRNAs of the cell has remained unknown.

In this study, we addressed how TOE1 achieves specificity towards canonical Pol II snRNAs. Global nascent small RNA 3’ end sequencing in the absence or presence of TOE1 confirms that TOE1 specifically processes canonical Pol II snRNAs over other classes of small RNAs. Dissecting the features of Pol II snRNPs recognized by TOE1 revealed the Sm complex and the TMG cap as two key features characteristic of Pol II snRNPs that promote TOE1 processing. An unstable U1 snRNA variant known to escape TOE1-processing, U1v15, is fully processed by TOE1 when the variant Sm binding motif is converted into that of canonical snRNAs. Our findings demonstrate that TOE1-mediated snRNA maturation is driven by Sm complex assembly and cap trimethylation, features that are specific to canonical Pol II snRNAs undergoing proper biogenesis.

## Results

### TOE1 specifically processes Pol II snRNAs

RNAs affected at steady state by TOE1 depletion have been previously globally monitored^11^. Since the processing of stable RNAs is better captured in the nascent RNA population, to further delineate the repertoire of small RNA targets of TOE1, we isolated nascently transcribed small RNAs, 100 to 500 nucleotides in length, from human embryonic kidney 293T (HEK293T) cells under normal or TOE1-depleted conditions and subjected them to global 3’ end sequencing (Figure 1a and Supplementary Figures S1a and S1b). Comparing mean genomic-encoded tail lengths of small RNAs in the absence or presence of TOE1 revealed Pol II snRNAs as the primary targets of TOE1-mediated 3’ end processing (Figures 1b and 1c). A subset of Pol II snRNAs also accumulated short oligo(A) tails in the absence of TOE1 (Figures 1d and 1e). Only one small ncRNA that was not an snRNA, scaRNA20, was observed to be significantly extended in the absence of TOE1 (Figures 1b and Supplementary Figure S1c), primarily due to an extended oligo(A) tail (Supplementary Figure S1d). While our global sequencing assays captured the majority of Pol II snRNAs as targets of TOE1, U5 and U4atac snRNAs were of too low abundance in the sequencing assays to include in the analysis; however, these have both previously been identified as TOE1 targets in gene-specific sequencing experiments^10,12^. Collectively, our nascent global small RNA 3’ end sequencing data show TOE1 specificity towards Pol II snRNAs as a class, although TOE1 may also act on a subset of other small ncRNAs, including scaRNA20, consistent with previous reports^11,26^. This raised the question of how TOE1 achieves specificity for Pol II snRNAs over other small ncRNAs.

**Figure 1.**
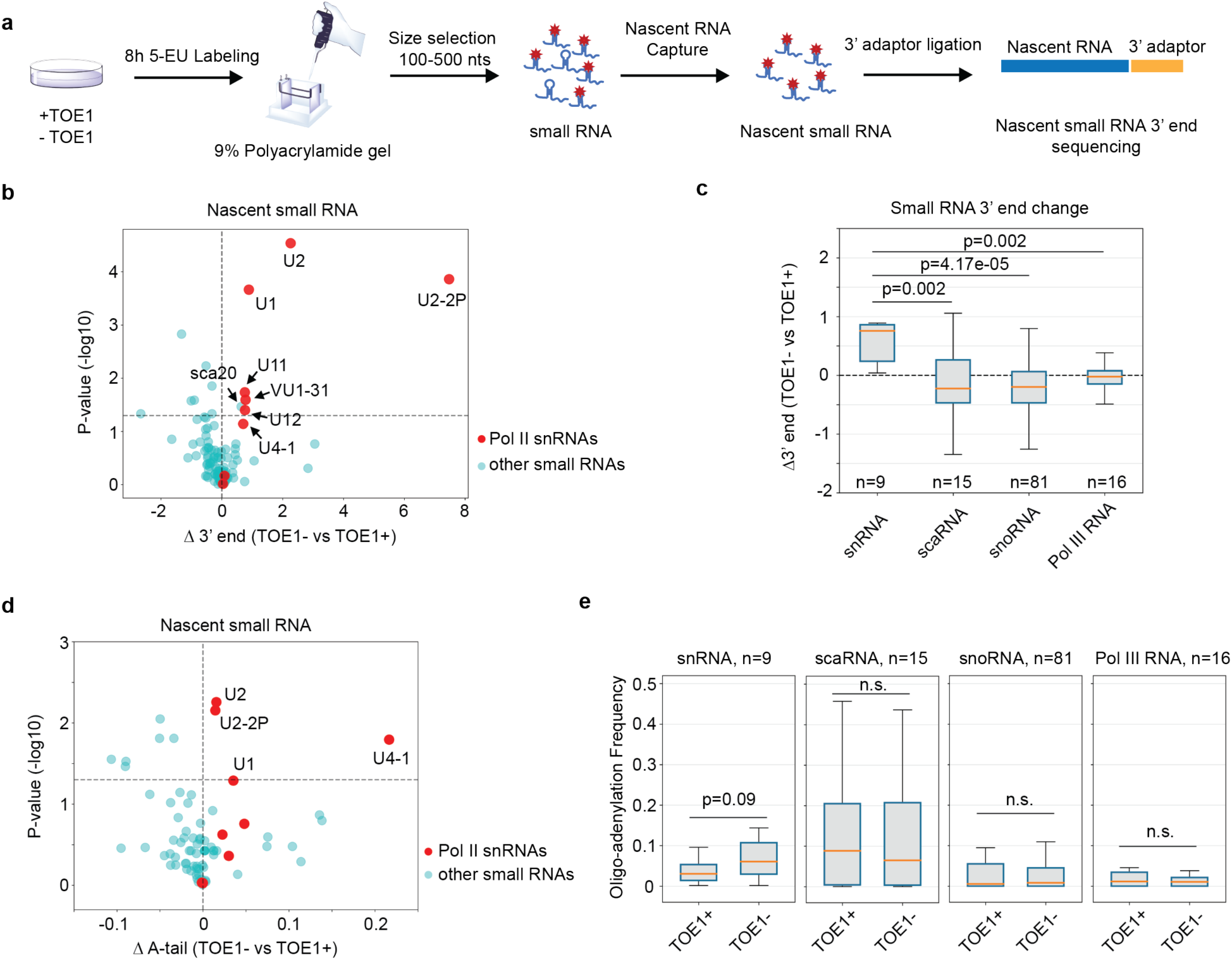
TOE1 shows specificity towards Pol II snRNAs. (**a**) Schematic of the nascent small RNA global 3’ end sequencing workflow. Control (TOE1+) or TOE1 depleted (TOE1-) cells were metabolically labelled with 5-ethynyl uridine (5-EU) followed by size selection and nascent RNA capture. Nascent RNA 3’ ends were determined by global small RNA sequencing. **(b)** Scatter plot showing changes in nascent small RNA 3’ end lengths in TOE1-versus TOE1+ cells and corresponding p-values. Each dot represents an individual small ncRNA with Pol II snRNAs in red and other small ncRNAs in cyan. P-values were calculated from three individual biological repeats by a two-tailed Student’s t-test and plotted on a –log10 scale. The horizontal dashed line represents p=0.05. **(c)** Box and whisker plots showing changes in 3’ end lengths of groups of related small ncRNAs in TOE1-versus TOE1+ cells. The number (n) of RNAs in each group is indicated. Boxes represent quartiles around median values shown as orange lines, with whiskers extending to maximum and minimum values within 1.5 times the interquartile range from the boxes. P-values were calculated by comparing each group of small ncRNAs to the group of Pol II snRNAs using two-sample Kolmogorov-Smirnov test. **(d)** Scatter plot of changes in frequencies of nascent small RNAs containing 3’ end posttranscriptional oligo(A) tails, defined as two or more post-transcriptional adenosines, in TOE1-versus TOE1+ cells. Dots and axes are labelled as in panel *b*. **(e)** Box and whisker plots showing distributions of 3’ end oligoadenylation (two or more adenosines) frequencies for each group of small ncRNAs in TOE1+ or TOE1-cells. Shown p-values were calculated using Student’s two-tailed t-test comparing TOE1-to TOE1+ conditions.

### Mutations in Sm complex and U1-70K binding motifs of U1 snRNA impair TOE1-mediated 3’ end processing

To investigate how TOE1 achieves substrate specificity, we turned to U1 snRNA (Figure 2a). Reasoning that TOE1 may recognize specific protein components of the U1 snRNP, we introduced previously described mutations that disrupt the interaction of U1 snRNA with U1A, U1-70K, and the Sm complex (Figure 2b)^47,58,59^. These mutations were introduced into a bar-coded exogenous U1 snRNA^10,57^, which allowed us to monitor their effect on U1 snRNA processing by 3’ end sequencing. Consistent with early observations in Xenopus oocytes^56^, disruption of the Sm complex binding motif led to a strong defect in U1 snRNA 3’ end processing (Figures 2c and 2d). In addition, mutating the U1-70K binding motif caused a minor, but statistically significant, defect in U1 snRNA 3’ end processing, whereas disrupting the U1A binding motif did not impair processing (Figures 2c and 2d). Further analyses of the mutant U1 snRNA 3’ ends revealed that the disruption of the Sm complex binding motif caused a significant fraction of the U1 snRNA population to accumulate with oligo(U) tails (Figures 2e and 2f), consistent with recent observations^19^. Interestingly, U1 snRNA mutated for U1-70K binding also accumulated oligo(U) tails, and both Sm and U1-70K binding mutants showed oligo(A) tailing as well (Figures 2e and 2g). Monitoring 3’ end processing of the mutant U1 snRNAs in the presence or absence of TOE1 depletion revealed that disruption of the Sm complex binding motif fully abolished the ability of TOE1 to process U1 snRNA (Figures 2h and 2i), whereas processing by TOE1 is only partially defective for the U1-70K binding mutant (Supplementary Figures 2a and 2b).

**Figure 2.**
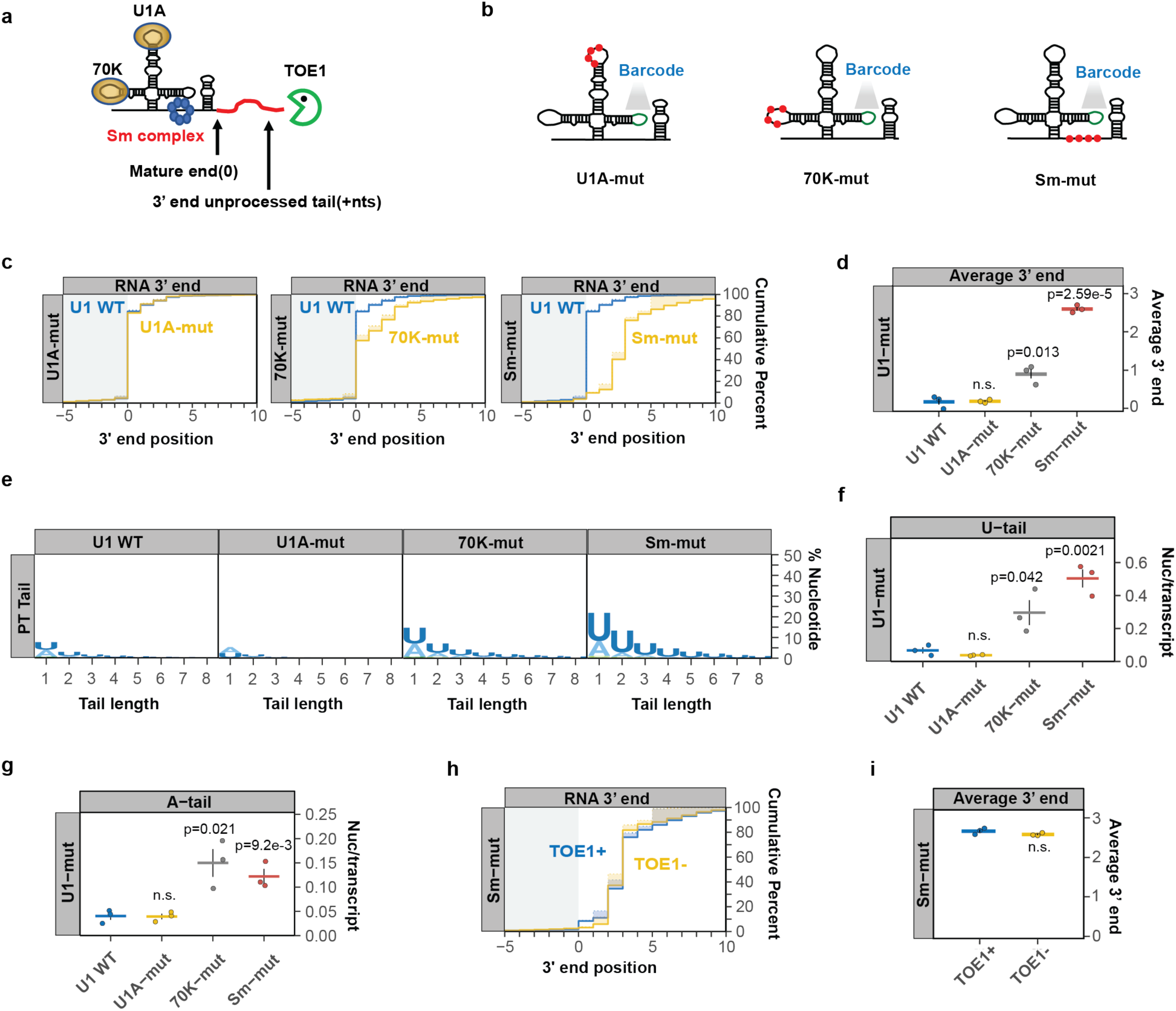
Mutations in Sm complex– and U1-70K-binding motifs of U1 snRNA inhibit 3’ end processing. (**a**) Schematic of the U1 snRNP with 70K, U1A, and Sm complex components shown at their binding motifs and a red line representing a 3’ end unprocessed tail. **(b)** Schematics of barcoded U1 snRNAs with mutations known to abolish binding of U1A (U1A-mut), U1-70K (70K-mut) or the Sm complex (Sm-mut) (Supplementary TableS2) shown as red dots. A barcode sequence in stem loop 3 is indicted in green. **(c)** Cumulative plots showing distributions of 3’ end positions for each U1 snRNA mutant (yellow) as compared to wild type U1 snRNA (U1 WT, blue) as determined by U1 snRNA gene-specific 3’ end sequencing. Mature 3’ ends are represented as position 0 with positive numbers representing unprocessed 3’ end tails. Solid lines represent actual snRNA 3’ end positions, while dotted lines represent genome-encoded 3’ end positions with shading in between representing post-transcriptionally added nucleotides. Only RNAs terminating at or downstream of the –10 position were analyzed. Three individual biological repeats were averaged for each plot. **(d)** Average 3’ end positions for each U1 WT or mutant snRNAs with each dot representing an individual biological repeat. Black vertical lines represent standard error of means (SEM) and p-values were calculated using Student’s two-tailed t-test comparing U1 mutants to U1 WT (p> 0.05 is noted as n.s.). **(e)** Sequence logo plots representing percentages of post-transcriptional added nucleotides for U1 WT and U1 mutant snRNAs. **(f, g)** Average post-transcriptional added uridines (*f*) and adenosines (*g*) per transcript for U1 WT and U1 mutants plotted as in panel *d*. **(h)** Cumulative plot showing 3’ end distributions for the Sm mutant U1 snRNA under TOE1+ (blue) or TOE1-(yellow) conditions. **(i)** Average 3’ end positions for the Sm mutant U1 snRNA in TOE1+ (blue) and TOE1-(yellow) conditions.

### Sm complex depletion impairs 3’ end processing of multiple Pol II snRNAs

Sm complex assembly is a feature characteristic of all Pol II snRNAs. To test if Sm complex assembly is necessary for 3’ end processing of Pol II snRNAs in general, we depleted the Sm complex component SmB (Supplementary Figure 3a) and performed targeted 3’ end sequencing of nascent Pol II snRNAs. All tested Pol II snRNAs showed defects in 3’ end processing following SmB depletion as compared to the negative control U3 snoRNA, although some snRNAs were affected more than others (Figure 3a and Supplementary Figure S3). U1 and U2 snRNAs showed strong defects in 3’ end processing, whereas U4, U5, and U4atac snRNAs showed significant, but less extensive, defects. A majority of the Pol II snRNAs also accumulated oligo(U) tails to various degrees upon SmB depletion (Figures 3b and 3c). These tails were most prominent for U1, U2 and U5 snRNAs, and observed at much lower levels for U4 and U4atac snRNAs (Figures 3b, 3c and Supplementary Figure S3). The control U3 snoRNA showed a very minor increase in uridylation upon SmB depletion (Figures 3b and 3c). Collectively, our findings demonstrate a general role for the Sm complex in 3’ end processing of Pol II snRNAs, although some snRNAs may be more sensitive to Sm complex levels than others.

**Figure 3.**
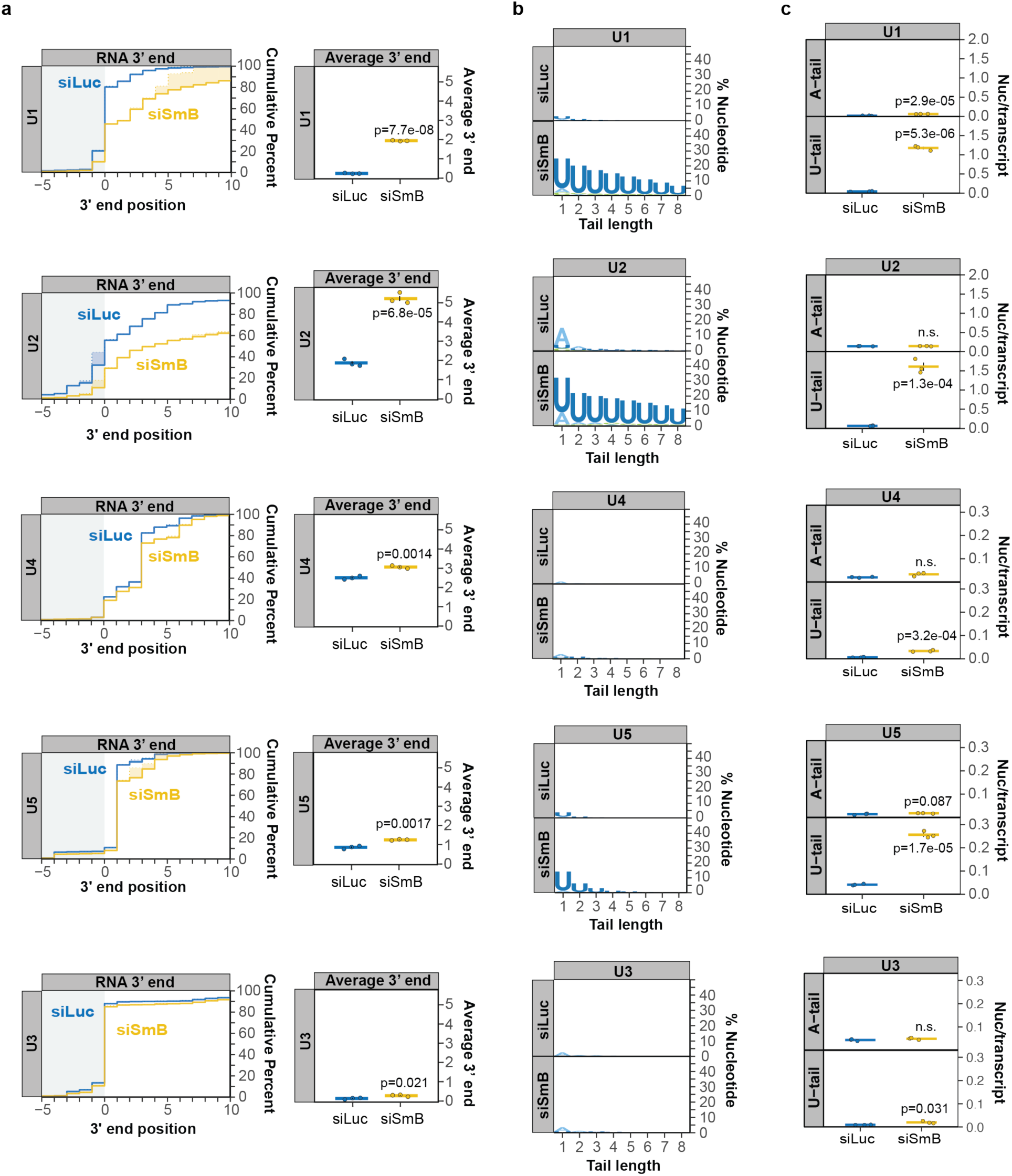
Depletion of Sm complex component SmB impairs 3’ end processing of multiple Pol II snRNAs. (**a**) Cumulative plots showing 3’ end distributions of nascent U1, U2, U4, and U5 snRNAs in control (siLuc; blue) versus SmB (siSmB; yellow) knock-down conditions. U3 snoRNA serves as a negative control. The average of three individual biological repeats are plotted and average 3’ end positions are graphed on the right with dots representing individual biological repeats and SEMs shown as vertical black lines. P-values were calculated by Student’s two-tailed t-test. **(b)** Sequence logo plots representing the percentage of post-transcriptionally added nucleotides for snRNAs and U3 snoRNA under control (siLuc) or SmB knock down (siSmB) conditions. **(c)** Average number of post-transcriptionally added adenosine or uridine nucleotides per transcript for major class snRNAs and U3 snoRNA plotted as in panel *a*.

### U1-70K stimulates U1 snRNA 3’ end processing by promoting Sm complex assembly

In contrast to the Sm complex, which assembles with all Pol II snRNAs, U1-70K is unique to U1 snRNA. Previous studies have shown that U1-70K helps promote assembly of the Sm complex onto U1 snRNA, which contains a suboptimal Sm binding site^41^ (Figure 4a). Consistent with the effect of the U1-70K binding-site mutation, depletion of the U1-70K protein impaired nascent U1 snRNA 3’ end processing and caused low-level oligo(A) and oligo(U) tailing (Figures 4b-4e and Supplementary Figure S4a). To test whether U1-70K promotes U1 snRNA 3’ end processing via its stimulation of Sm complex assembly, we tested the effect of converting the suboptimal U1 snRNA Sm binding motif into a canonical one (Figure 4f, superU1), known to promote assembly of the Sm complex independently of U1-70K^41^. The canonical Sm binding site rescued the 3’ end processing defect of the U1-70K binding site mutation and strongly reduced adenylation and uridylation (Figures 4g-4i). Thus, U1-70K promotes U1 snRNA 3’ end processing by stimulating Sm complex assembly. In contrast to U1-70K and the Sm complex, depletion of U1 snRNP components U1A and U1C showed no defect in U1 snRNA 3’ end processing (Supplementary Figure S4b-g), consistent with our observations with the U1A-binding mutant U1 snRNA.

**Figure 4.**
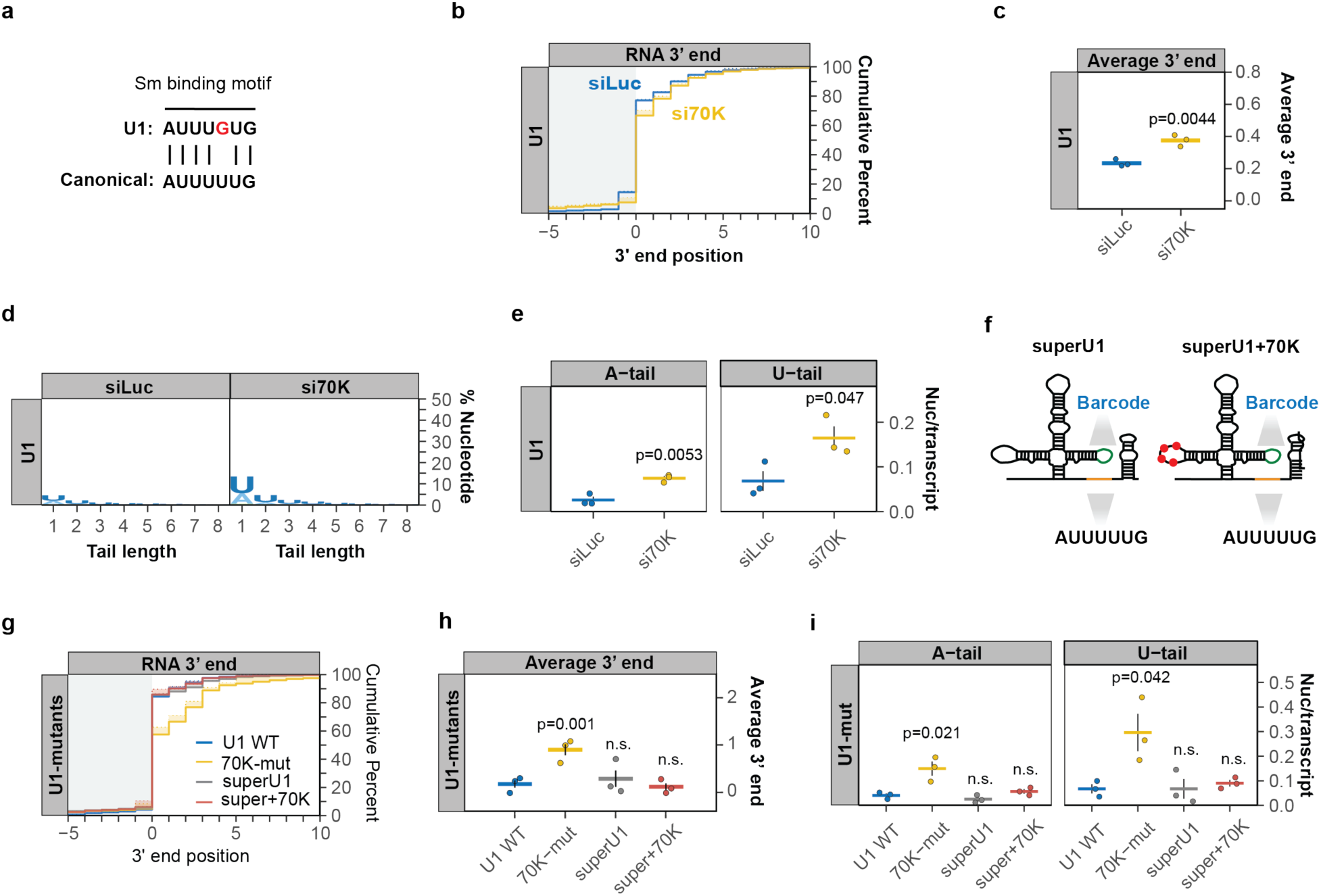
U1-70K stimulates U1 snRNA 3’ end processing by promoting Sm complex assembly. **(a)** Comparison of the U1 snRNA Sm binding motif with the canonical Sm binding motifs found in other Pol II snRNAs. **(b)** Cumulative plot showing the 3’ end distributions of nascent U1 snRNA in control (siLuc, blue) or U1-70K depletion (si70K, yellow) conditions. Averages of three individual biological repeats are plotted. **(c)** Average nascent U1 snRNA 3’ end positions from the experiments plotted in panel *b* with dots representing individual biological repeats and SEMs shown as vertical black lines. The p-value was calculated by Student’s two-tailed t-test. **(d)** Sequence logo plot representing the percentage of post-transcriptionally added nucleotides in control (siLuc) or U1-70K depleted (si70K) conditions. **(e)** Average post-transcriptional added adenosines and uridines per transcript plotted as in panel *c*. **(f)** Schematics of barcoded exogenous U1 snRNAs with canonical Sm binding motif (superU1) and superU1 with mutations that abolish U1-70K binding shown as red dots (superU1+70K). **(g)** Cumulative plot showing 3’ end distributions for indicated exogenous U1 snRNAs. **(h)** Average 3’ end positions for indicated exogenous U1 snRNAs, plotted as in panel *c*. **(i)** Average post-transcriptionally added adenosine and uridine nucleotides per transcript for indicated exogenous U1 snRNAs, plotted as in panel *e*.

### The U1 snRNA variant U1v15 escapes TOE1 processing due to variant nucleotides in its Sm-binding motif

The human U1 snRNA variant U1v15 is known to evade TOE1 recognition and undergo rapid degradation^12^. To identify the nucleotide variations within U1v15 RNA that inhibit processing by TOE1, we introduced variant nucleotides found within the U1A or Sm binding motifs of U1v15 into the bar-coded canonical U1 snRNA (Figure 5a). The variant nucleotides of the U1v15 U1A binding motif showed no effect on 3’ end processing when introduced into the canonical U1 snRNA (Figures 5b and 5c). By contrast, the variant nucleotides of the Sm binding motif were sufficient to trigger a 3’ end processing defect similar to that seen for the variant U1v15 snRNA (Figure 5b and 5c), and promote similar levels of oligo(U) and –(A) tailing (Figures 5d-5f). We next converted the variant Sm binding site of U1v15 RNA into the Sm binding motif found in U1 snRNA (Figure 5a, v15-SmWT) or the canonical Sm binding motif (Figure 5a, v15-super), while leaving all other variant nucleotides intact. These modifications rescued 3’ end processing of the U1v15 snRNA variant (Figures 5b and 5c), although a minor residual level of mono-adenylation and –uridylation could be observed (Figures 5d-5f). Taken together, these observations show that an intact Sm complex binding site is not only necessary for the processing of canonical Pol II snRNAs by TOE1, but can also rescue the processing of a normally unprocessed and unstable snRNA variant.

**Figure 5.**
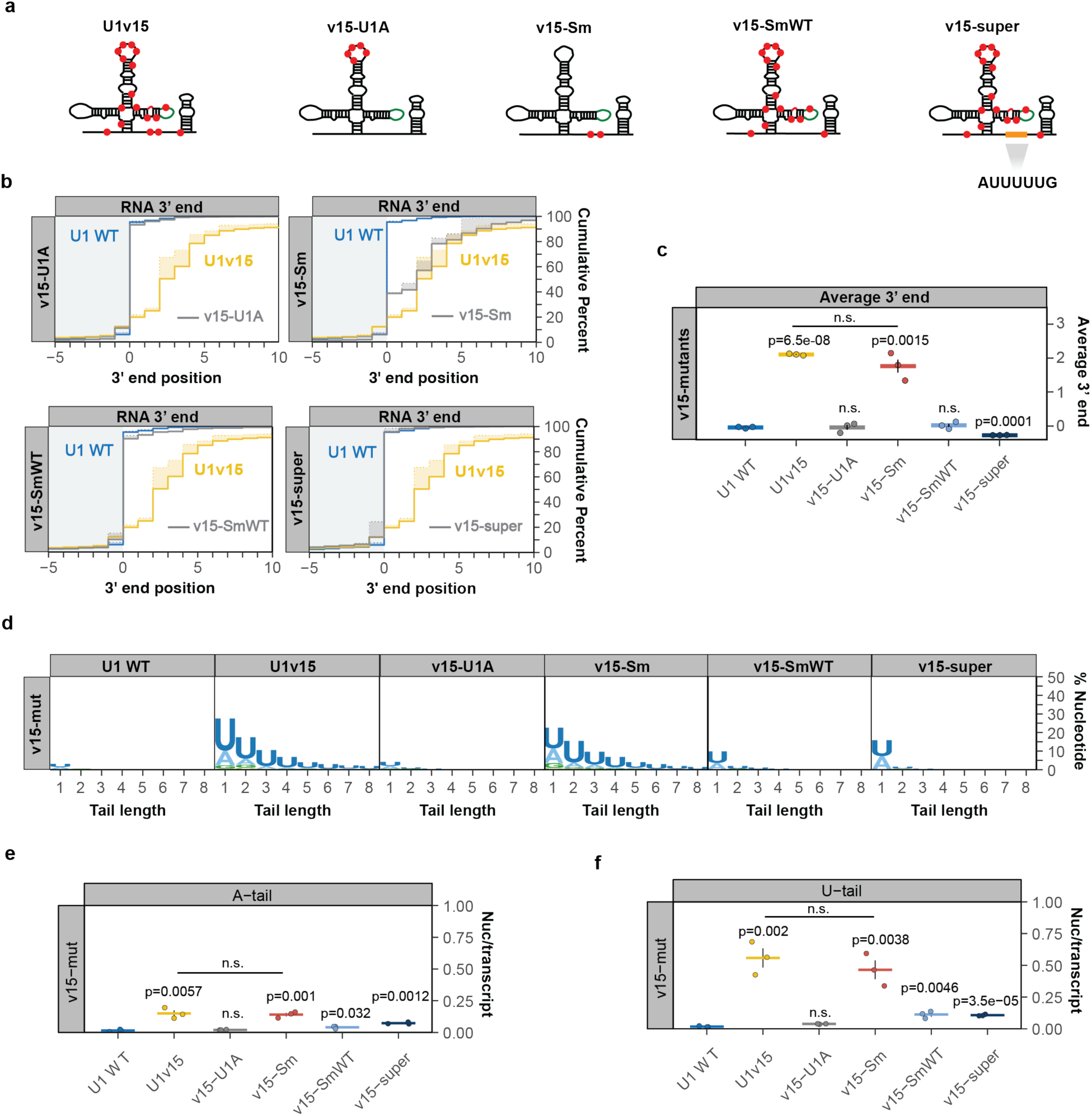
U1 snRNA variant U1v15 escapes TOE1 processing due to variant nucleotides in its Sm binding motif. (**a**) Schematic of U1 snRNA variant U1v15 and U1 snRNA mutants (v15-U1A, v15-Sm, v15-SmWT) (Supplementary TableS2) containing mutations corresponding to U1v15 nucleotide variations indicated by red dots. v15-super corresponds to U1v15 with a canonical Sm binding motif. **(b)** Cumulative plots showing 3’ end distributions for U1 WT (blue), U1v15 (yellow), and each v15 mutant (grey). The average of three individual biological repeats are plotted. **(c)** Average 3’ end positions for U1 WT, U1v15, and v15 mutants. Dots represent individual repeats and vertical lines are SEM. P-values were calculated by Students two-tailed t-test by comparing v15 mutants to U1 WT. **(d)** Sequence logo plots representing the percentage of posttranscriptional added nucleotides for U1 WT, U1v15, and v15 mutants. **(e, f)** Average post-transcriptionally added adenosines (*e*) and uridines (*f*) per transcript plotted as in panel *c*.

### TOE1 directly recognizes the Sm complex-assembled U1 snRNP

The assembly of the Sm complex onto Pol II snRNAs takes place in the cytoplasm and triggers subsequent 5’ cap trimethylation and nuclear import^50–54^ (Figure 6a). Since TOE1 is known to concentrate in the nucleus^11,20^, our findings raised the question of whether TOE1 directly recognizes the Sm complex-assembled snRNP. Alternatively, the Sm complex could promote TOE1-mediated processing by indirect means such as via snRNA nuclear import. To address this question, we tested the ability of recombinant TOE1 to process wild-type or Sm-mutant snRNPs isolated from cells. A 2’-OMe-RNA oligonucleotide (Supplementary Table S1) hybridizing to the U1 snRNA 5’ splice site recognition motif was used to isolate bar-coded U1 wild-type or Sm-mutant snRNP complexes exogenously expressed in TOE1-depleted HEK293T cells (Figure 6b and Supplementary Figure S5). Isolated snRNPs were subsequently incubated with Flag-tagged TOE1 protein *in vitro* and analyzed by 3’ end sequencing. We found that the wild-type U1 snRNP was increasingly processed at the 3’ end with increasing concentrations of TOE1, but not with a previously characterized catalytically inactive TOE1 mutant^10^ (Figures 6c-e). By contrast, the Sm-mutant U1 snRNP showed little processing by TOE1 (Figures 6f-h). These observations demonstrate that TOE1 directly recognizes the Sm complex-assembled U1 snRNP.

**Figure 6.**
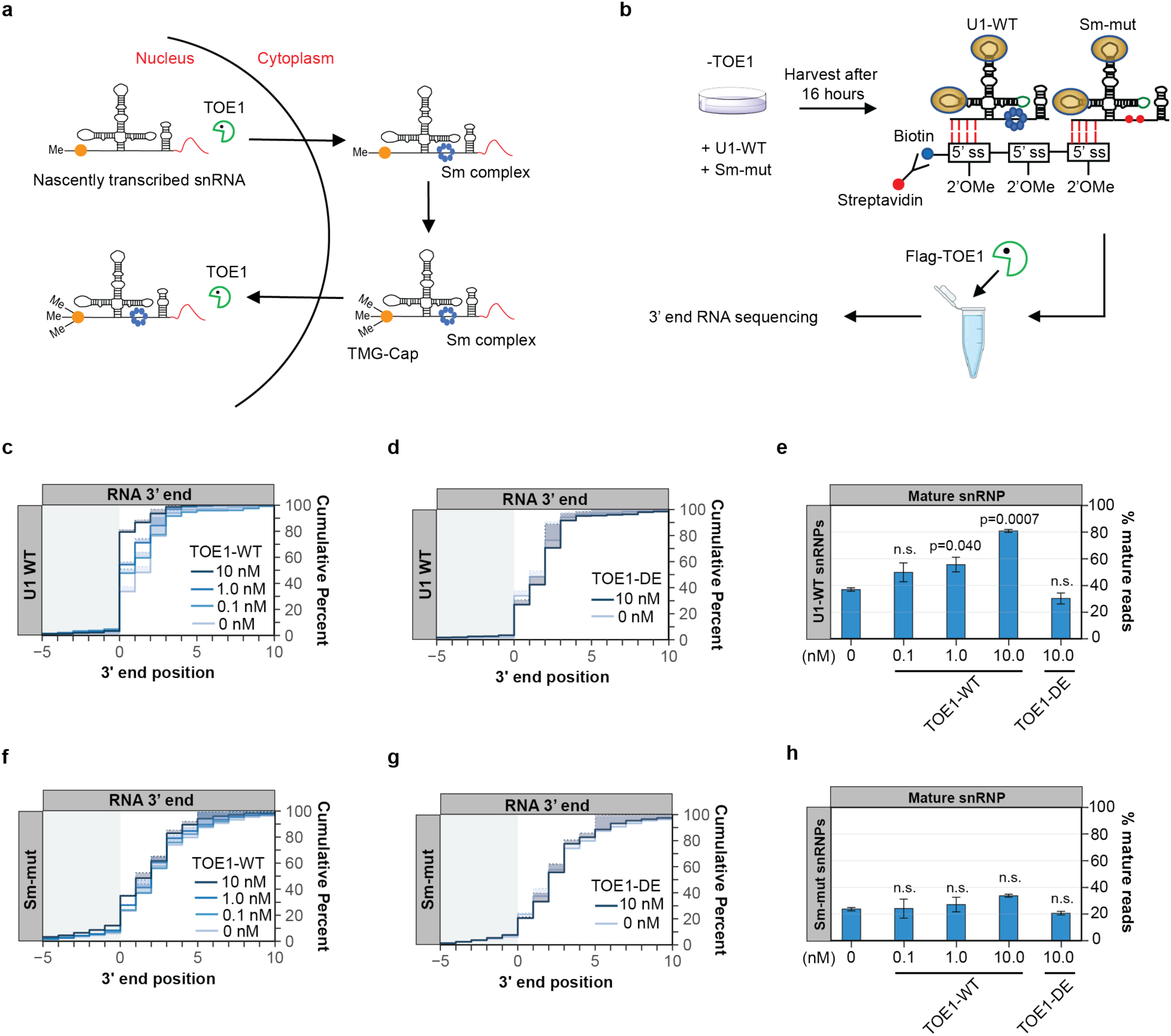
TOE1 preferentially processes wild-type over Sm-mutant U1 snRNP *in vitro*. (**a**) Schematic of the Pol II snRNA biogenesis pathway highlighting 3’ end adenylation, Sm complex assembly, 5’ cap trimethylation, and TOE1-mediated processing steps. **(b)** Schematic of the *in vitro* U1 snRNP pull-down and TOE1 processing assay. See methods section for details. **(c, d)** Cumulative plots showing 3’ end distributions of wild type U1 (U1 WT) snRNPs after incubation with TOE1 (panel *c*) or catalytically inactive TOE1 (TOE1 DE, panel *d*) at indicated concentrations. The average of two individual biological repeats are plotted. **(e)** Percentage of fully processed U1 WT snRNPs after TOE1 or TOE1 DE incubation at indicated concentrations. P-values were calculated by Student’s one-tailed t-test by comparing with the 0 nM TOE1 condition (n.s.: p>0.05). **(f-h)** Same as panels *c-e* but analyzing the U1 Sm-mut snRNP.

### snRNA processing by TOE1 is activated by the 5’ TMG cap

In addition to promoting nuclear import, Sm complex assembly also stimulates trimethylation of the snRNA cap^50–52^. Interestingly, the TOE1 homolog PARN is known to be activated by an m7G-monomethyl cap^60–62^. To test if TOE1 activity is affected by the snRNA cap structure, we generated uncapped, m7G-capped, or TMG-capped U1 snRNAs by *in vitro* transcription (Supplementary Figures S6a-c). Each U1 snRNA contained a 20-nucleotide genomic-encoded tail to allow for monitoring of TOE1 activity (U1+20, Figure 7a). Incubating each of the differently capped U1 snRNAs with TOE1, we found that TOE1 processed the 5’ TMG-capped U1 snRNA much more efficiently than the corresponding 5’ uncapped or m7G-capped U1 snRNA substrates (Figures 7b and 7c). The TMG cap analog used in this assay produces an RNA population of which at most half are TMG capped (Supplementary Figures S6a and S6b), which may explain why about half of the RNA population remaining unprocessed by TOE1 (Figure 7b). Collectively, our observations demonstrate that TOE1 achieves substrate specificity through the recognition of Sm complex assembly and cap trimethylation, two features that distinguish Pol II snRNAs undergoing proper assembly from other small ncRNAs of the cell.

**Figure 7.**
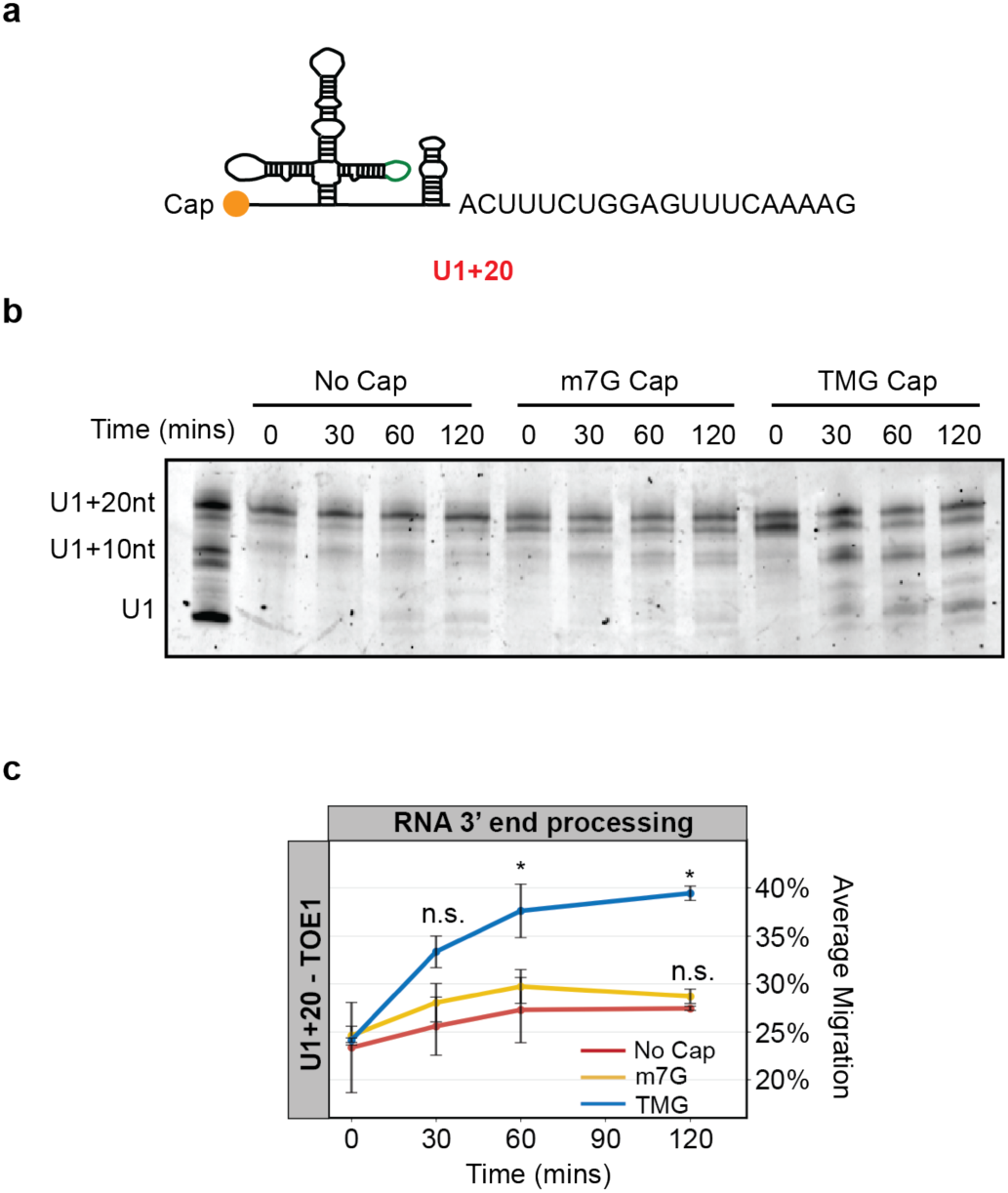
U1 snRNA processing by TOE1 is stimulated by the 5’ TMG cap. (**a**) Schematic of *in vitro* transcribed U1 snRNA with a 20-nucleotide (nt) genomic-encoded tail (U1+20). **(b)** Representative denaturing gel showing uncapped, m7G-, or TMG-capped U1+20 incubated with TOE1 *in vitro* for indicated times. The RNA ladder consists of U1 snRNA with no tail, a 10-nt, or a 20-nt tail. **(c)** Quantification of TOE1 processing in panel *b*. The average RNA migration for each condition was quantified by ImageJ (with the U1+20-nt marker corresponding to 0% migration and U1 corresponding to 100% migration). The average of two individual biological repeats is plotted. P-values were calculated by Student’s two-tailed t-test comparing m7G or TMG U1+20 migration to the uncapped U1+20 migration at each timepoint (*: p<0.05; n.s.: p>0.1).

## Discussion

Pontocerebellar Hypoplasia 7-associated protein TOE1 is a key 3’ end maturation factor for canonical Pol II snRNAs, but how it distinguishes substrate from non-substrate RNAs has remained unknown. Here, we present evidence that TOE1 selectively processes Pol II snRNAs over other classes of small non-coding RNAs (Figure 1). This specificity is imparted in part by Sm complex assembly, a distinguishing feature of Pol II snRNAs. Indeed, manipulations known to inhibit Sm complex assembly, including U1 snRNA mutations and nucleotide variations (Figures 2 and 5), and depletion of an Sm complex component or U1-70K (Figures 3 and 4), disrupt TOE1-mediated snRNA 3’ end processing. TOE1 directly recognizes the Sm complex-assembled snRNP as demonstrated by its specificity towards wild-type over Sm-mutant U1 snRNP in an *in vitro* 3’ end processing assay (Figure 6). TOE1 also recognizes the trimethylated snRNA cap as evidenced by the specificity of TOE1 for a 5’ TMG capped snRNA *in vitro* (Figure 7). These findings reveal the molecular basis for how TOE1 distinguishes canonical Pol-II snRNAs from other small ncRNAs of the cell.

When and where during their maturation does TOE1 process snRNAs? The biogenesis of Pol II snRNAs involves both nuclear and cytoplasmic processes (Figure 6a). We previously presented evidence that TOE1 acts on snRNAs at least twice during their biogenesis, by first partially processing snRNAs prior to or during nuclear export, before completing maturation during or after nuclear re-import^12^. Our findings presented here provide a molecular basis for how TOE1 achieves specificity towards snRNAs during late-stage biogenesis. At this point, snRNAs have undergone Sm complex assembly and cap trimethylation, the two features shown here to promote processing by TOE1. Since snRNA cap trimethylation occurs as a consequence of Sm complex assembly^50–52^, it is possible that this late-stage processing by TOE1 occurs solely through TOE1 recognition of the trimethylated cap. Consistent with a key role for the TMG cap, depletion of TGS1, the enzyme that carries out cap trimethylation, has been reported to lead to accumulation of 3’ end extended snRNAs^63,64^. Alternatively, TOE1 may additionally directly recognize the Sm complex. The early-stage processing of snRNAs by TOE1, which occurs prior to Sm complex assembly and cap trimethylation, is likely a consequence of the activity of TOE1 as a deadenylase^10–12,20^. Consistent with this idea, TOE1 can process a U1 snRNA containing an oligo(A) tail to maturity *in vitro*, independent of its cap structure (Supplementary Figures S6d-f).

Hundreds of unstable snRNA variants are encoded in the human genome^65,66^. We previously observed that TOE1 distinguishes canonical snRNAs from tested unstable snRNA variants, leaving the latter unprocessed^12^. Our observations presented here demonstrate that U1v15 RNA escapes TOE1 recognition specifically due to nucleotide variations in the Sm binding motif (Figure 5). These same nucleotide variations also trigger 3’ end oligo(U) and –(A) tailing (Figure 5), features known to promote degradation by exonucleases and the decapping machinery^17–19,57^. Thus, multiple layers of quality control act on U1v15 RNA to prevent it from assembling into spliceosomes. This likely represents general mechanisms that serves to repress the subset of transcribed snRNA variants that have acquired nucleotide variations that impair Sm complex assembly, either in the Sm binding motif or in other regions of the snRNA that affect Sm binding, such as the U1-70K binding site (Figures 2 and 4).

A large number of small non-coding RNAs are processed by enzymes of the DEDD deadenylase family, including TOE1 and PARN^10,11^. Despite their homology and shared preference for oligo(A) tails, TOE1 and PARN show different specificities for small RNA substrates^10–12^. How do these two enzymes achieve specificity towards different substrates? Our findings suggest that this may be explained for a subset of target RNAs by their cap structures. While we find here that TOE1 recognizes the TMG cap characteristic of Pol II snRNAs, PARN was previously shown to be activated by the m7G cap^60–62^, a 5’ modification found on a subset of small ncRNAs targeted by PARN, including TERC and a subset of snoRNAs. There are reports that TERC can also be observed with a TMG cap^67^ and serve as a substrate for TOE1^26^, although we have been unable to detect processing of TERC by TOE1 in HEK293T cells^10^. While cap structures may explain the differential activity of TOE1 and PARN towards a subset of substrates, how PARN achieves specificity for other substrates, including a majority of snoRNAs that are processed from introns and therefore remain uncapped^68^, remains unresolved. A contributing factor may well be subcellular localization, with TOE1 known to co-localize with snRNAs in Cajal Bodies and PARN with snoRNAs and other substrates in the nucleolus. Similarly, the molecular basis for target preferences of other maturation exonucleases PNLDC1 and USB1 remain an important question for future study.

The 3’ ends of nascent small non-coding RNAs serve as quality control hubs for competing exonucleases to drive 3’ end maturation or degradation. We previously identified TOE1 and the nuclear exosome as competing exonucleases acting on snRNA 3’ ends to distinguish canonical snRNAs from unstable snRNA variants^12^. Our findings here, that TOE1 recognizes late snRNA biogenesis features of Sm complex assembly and cap trimethylation, reveals how TOE1 can achieve specificity towards canonical snRNPs undergoing correct assembly. Unstable snRNA variants^12,66^ and canonical snRNAs experiencing defects in snRNP assembly^19,57^, likely undergo degradation because they fail to reach this late stage of snRNP biogenesis. This may represent a general principle in RNA quality control whereby substrate specificity relies on maturation enzymes that specifically recognizes canonical RNAs undergoing proper biogenesis competing with more promiscuous degradation enzymes to distinguish normal from aberrant RNAs and ultimately dictate their fate.

## Data accessibility

RNA sequencing data have been deposited to the Gene Expression Omnibus (GEO) under accession number GSE240774.

## Acknowledgment

We thank Tim Nicholson-Shaw for sequencing analysis support. We thank Alberto Carreño, Tim Nicholson-Shaw, Cody Ocheltree, Megan Dowdle, and Stephen Sanders for useful discussion of the manuscript. We acknowledge the UCSD Stem Cell Genomics and Microscopy Core for use of their Illumina Miseq instrument. This work was supported by NIH grant R35 GM118069 to J.L-A. R.M.L. was the recipient of a National Research Service Award Postdoctoral Fellowship (NIH F32 GM106706) and was a San Diego IRACDA Fellow (NIH K12 GM06852). This publication includes data generated at the UC San Diego IGM Genomics Center utilizing an Illumina NovaSeq 6000 that was purchased with funding from a National Institutes of Health SIG grant (#S10 OD026929).

## Author Contributions

T.M., E.S.X., and R.M.L. performed all experiments. T.M., E.S.X., R.M.L., and J.L-A. performed data analyses. T.M. and J.L-A. wrote the manuscript.

## Methods

### Cell culture, RNA interference, and plasmid transfections

All cells were maintained in Dulbecco’s Modified Eagle Medium (DMEM, Gibco) supplemented with 10% Fetal Bovine Serum (FBS, Gibco) and 1% penicillin/streptomycin (Gibco) at 37°C, 5% CO_2_. Mycoplasma testing was routinely performed. TOE1 was depleted in HEK 293T-REx-derived TOE1-degron cells^12^ by incubation with 600 μM auxin hormone Indole-3-Acetic Acid (IAA, Sigma) for 8 hours. RNA interference was performed in HEK 293T Flp-In cells (FIRT, Thermo Fisher) with 20 nM small interfering (si)RNA targeting genes of interest, or luciferase as a control (Supplementary Table S1), using siLentFect (Bio-Rad) transfection reagent according to manufacturer’s recommendations at 72 and 24 hours before cell harvest. Plasmid transfections were performed using 2 μg plasmid per 3.5-cm well plates using Lipofectamine 2000 (Life Technologies) transfection reagent according to the manufacturer’s recommendations at 48 hours before harvest, unless specified otherwise.

### Nascent global small RNA 3’ end sequencing

HEK 293T-REx TOE1-degron cells were treated to deplete TOE1 as described above. 1.2 × 10^6^ TOE1-depleted, or control non-depleted, cells were incubated with 0.2 mM Ethynyl Uridine (EU; Thermo Fisher) for 8 hours and harvested in 1 ml TRIzol reagent (Thermo Fisher). Total RNA was isolated according to the manufacturer’s recommendation. Small RNA from 35 μg of total RNA from each sample was isolated by separation in a 9% polyacrylamide/6M urea denaturing gel. After Sybr Gold staining (Thermo Fisher), small RNAs between 100 and 500 nucleotides in length were cut out of the gel and eluted into 400 μl gel elution buffer (0.3 M sodium acetate pH 5.3, 1 mM EDTA, 0.1% SDS) by end-over-end rotation overnight at 4 °C. Eluted small RNAs were subsequently subjected to phenol:chloroform:isoamyl alcohol extraction followed by ethanol precipitation as previously described^10^. Genomic DNA was removed using Turbo DNA-free kit (Thermo Fisher) and ribosomal (r)RNAs were depleted using RiboCOP rRNA depletion kit (Lexogen) per manufacturer’s recommendations. The rRNA-depleted small RNA samples were subsequently subjected to FastAP/PNK treatment to remove RNA 5’ and 3’ phosphates by incubating samples with 2.5 μl 10x FastAP Buffer (Thermo Fisher), 0.5 μl RNaseOUT (Thermo Fisher), 2.5 μl FastAP phosphatase (Thermo Fisher) in 25 μl total volume at 37°C for 30 minutes, followed by supplementation with 10 μl 10X PNK buffer (NEB), 1 μl 0.1 M DTT, 1 μl Turbo DNase (2 U/µl; Thermo Fisher), 1 μl RNaseOUT (40 U/µl; Thermo Fisher), and 7 μl Polynucleotide Kinase (10 U/µl; NEB) in a total of 100 μl and incubation for 20 minutes at 37 °C. RNA samples were purified using an RNA Clean & Concentrator kit (Zymo Research). AG10N or AG11N RNA adaptors (10 µM; Supplementary Table S1) were ligated to the RNA 3’ ends of each sample by incubation at 25°C for 75 minutes in a 20 µl total volume containing 1.8 μl DMSO (Sigma), 2 μl 10x T4 ligase buffer (NEB), 0.2 μl 0.1 M ATP (Thermo Fisher), 0.2 μl RNaseOUT (40 U/µl; Thermo Fisher), 8 μl 50% PEG 8000 (NEB), 1.3 μl T4 RNA ligase (30 U/µl; NEB). RNA samples were then subjected to nascent RNA extraction using Click-it nascent RNA capture kit (Thermo Fisher) as previously described^12^. Reverse transcription was performed on the RNA capture beads in a total of 20 µl with 0.5 μl 20 μM AR17 primer (Supplementary Table S1), 2 μl 10x AffinityScript buffer (Agilent), 2 μl 0.1 M DTT, 0.8 μl 100 mM dNTPs, 0.3 μl RNaseOUT (40 U/µl; Thermo Fisher), 0.9 μl AffinityScript Reverse Transcriptase (Agilent) at 55°C for 45 minutes, followed by 15 minutes incubation at 70°C and 5 minutes of incubation at 85°C to release cDNA. Excess primers and RNA was removed from the first-strand cDNA by incubating with 3.5 μl ExoSAP-IT (Thermo Fisher) at 37°C for 15 minutes followed by addition of 3 μl of 1M NaOH at 70°C for 12 minutes. 3 μl of 1M HCl was added to the sample after the clean-up to adjust pH. A 3Tr3 adaptor (Supplementary Table S1) was ligated to cDNA 5’ ends by adding 0.8 μl of the adaptor at 80 µM to the cDNA sample along with 1 μl 100% DMSO (Sigma), 2 μl 10×T4 ligase buffer (NEB), 0.2 μl 0.1 M ATP (Thermo Fisher), 1.5 μl T4 RNA ligase (30 U/µl; NEB), and 1.1 μl double distilled water, and incubating the reaction at 25°C for 16 hours. The cDNA was amplified in two stages of Polymerase Chain Reaction (PCR) using Q5 DNA polymerase (NEB). For the first PCR reaction, the cDNA library was amplified using 3’ adaptor primer (AR17) and a primer complementary to the 5’ adaptor (RC_3Tr3) (Supplementary Table S1) for 8 cycles. The PCR product was purified by AMPure XP beads (Beckman Coulter) per manufacturer’s recommendation. The second PCR reaction was performed using Illumina Trueseq D50X and D70X primers (Supplementary Table S1) for 18 cycles. Amplified cDNA was subsequently isolated by separation in a 3% agarose gel and isolation of cDNAs 200 to 600 base pairs in length (corresponding to 100 to 500 nucleotide RNAs plus adaptors) using QIAquick gel extraction kit (Qiagen) according to manufacturer’s recommendation. The library quality was monitored by qPCR for select genes and Tapestation (Agilent) analyses. 100 bp paired-end sequencing was performed on an Illumina Novaseq 6000 platform per manufaturer’s recommendation.

### Sequencing data analyses

FASTQ files were first subjected to 3’ adaptor and PCR duplicate removal using custom python scripts (https://pypi.org/project/jla-demultiplexer/). For highly abundant small RNAs (U1, U2, U3, 7SL and 7SK), 100,000 reads were selected for subsequent analyses by averaging three random samplings using seqtk (https://github.com/lh3/seqtk). This was done to prevent undercounting of these RNAs since their abundance in the full library exceeded the depth of the randomer used for detecting PCR duplicates (Supplementary Figure S1b). For all other RNAs, the entire library was used. Reads were mapped to the human genome (version hg38) using STAR 2.7.8a^69^. To diminish mis-mapping of canonical small RNA reads to small RNA variant genes, a three-pass alignment was applied. Briefly, reads were first mapped to the human hg38 genome (STAR –-outFilterMultimapNmax 1000 –-alignIntronMin 9999999 –-outFilterMultimapScoreRange –-outFilterMismatchNoverLmax 0.2) and reads mapping to small RNA genes were extracted using bedtools^70^ and samtools^71^. The mapped small RNA reads were then aligned to a custom FASTA database of canonical small RNA genes, each including 50 base pair upstream and downstream sequences (STAR –-outFilterMultimapNmax 1000 –-outFilterMultimapScoreRange 0 –-outFilterMismatchNoverLmax 0.2 –-outFilterMismatchNoverReadLmax 0.05 –-clip5pNbases 200--clip3pNbases 0 20 –-alignIntronMin 9999999 –-alignMatesGapMax 500 –-alignEndsType EndToEnd –-outReadsUnmapped Fastx). Reads that mapped to the canonical small RNA gene database were subsequently re-aligned to the full human h38 genome (STAR –-outFilterMultimapNmax 1000 –-outFilterMultimapScoreRange 0 –outFilterMismatchNoverLmax 0.025 –-alignIntronMin 9999999 –-alignMatesGapMax 500 –-alignEndsType Local) and those that again mapped to canonical small RNA genes were extracted with bedtools. This step was performed to remove canonical small RNA reads with sequencing errors that may misalign with small RNA variant genes. Reads that failed to map to the canonical small RNA gene database were also re-aligned with the full human hg38 genome using the same settings and combined with the reads mapping to canonical small RNA genes. Gene-specific 3’ end information and graphs were subsequently generated using Tailer^72^ (https://github.com/TimNicholsonShaw/tailer) using the global alignment mode.

### RNA gene-specific 3’ end sequencing

RNA was isolated using TRIzol (Thermo Fisher) per manufacturer’s recommendation. RNA was subsequently treated with DNase I (1 U/µl; Zymo research) in DNase buffer (10 mM Tris-HCl pH 7.5, 2.5 mM MgCl_2_, 0.5 mM CaCl_2_) at 25°C for 15 minutes, followed by extraction with phenol:chloroform:isoamyl alcohol and ethanol precipitation as previously described^12^. For nascent RNA sequencing, nascent RNA was captured as previously described^12^. Gene-specific RNA 3’ end sequencing libraries were generated using gene-specific primers (Supplementary Table S1) and sequenced on an Illumina MiSeq platform as previously described^12^.

### Isolation of Flag-tagged TOE1

HEK 293T Flp-In cells (FIRT) cells were transfected with pcDNA5-Flag-TOE1 plasmid^20^ as described above. 1 μg/ml of tetracycline was added 24 hours before harvest to induce Flag-TOE1 expression. 2 × 10^7^ Flag-TOE1-induced cells were lysed in 2.5 ml of isotonic lysis buffer (10 mM Tris-HCI pH 7.5, 150 mM NaCI, 2 mM EDTA, 0.1% Triton-X100, 1 mM PMSF, 2 µg/ml Aprotinin, 2 µg/ml Leupeptin) with 125 μg/ml RNase A. The lysate concentration was measured using BCA protein assay (Thermo Fisher) following manufacturer’s protocol. 40 µl of 50% anti-FLAG M2 agarose slurry (Sigma) was washed twice with 500 μl NET-2 (10 mM Tris HCl pH 7.4, 150 mM NaCl, 0.1% Triton-X100) before incubation with 1-5 mg of cell lysate with end-to-end rotation for 2 hours at 4°C. Beads were subsequently washed with 500 µl NET-2 eight times. Flag-TOE1 was eluted with 100 µl of NET-2 containing 150 ng/µl FLAG peptide (ApexBio) and 10% glycerol by rotating at 4°C for 30 minutes. The isolated protein was detected by silver staining (Thermo Fisher) per manufacturer’s recommendation. Eluates were aliquoted and stored at –80°C until use.

### In vitro snRNP pull-down and 3’ end processing assay

HEK293T-Rex TOE1-degron cells were treated to deplete TOE1 as described above and co-transfected with 0.2 μg U1-WT and 2 μg Sm-mutant (Supplementary Table S2) expression plasmids as described above, 16 hours prior to cell harvest. 2 × 10^7^ cells were suspended in 0.2 ml of low salt lysis buffer (10 mM Tris-HCI pH 7.5, 60 mM NaCI, 2 mM EDTA, 0.1% Triton-X100, 1 mM PMSF, 2 µg/ml Aprotinin, 2 µg/ml Leupeptin, 0.1 µg/ml yeast total RNA, 2 U/ml RNaseOUT) and incubated on ice for 10 minutes. 20 µl of 0.5 µM 2’OMe-RNA-oligo probe (Supplementary Table S1), 12 µl streptavidin magnetic beads (NEB), and 150 µl wash/binding buffer (0.5 M NaCl, 20 mM Tris-HCl pH 7.5, 1 mM EDTA) were incubated on ice for 30 minutes with occasional agitation. 200-400 µl of cell lysate was added to each tube and incubated at 4°C for 2 hours.

Beads were washed twice with 100 µl wash/binding buffer followed by one wash with 100 µl low salt wash buffer (20 mM Tris-HCl pH 7.5, 150 mM NaCl, 1 mM EDTA), and 5 times with 100 µl EDTA-free low salt buffer (20 mM Tris-HCl pH 7.5, 150 mM NaCl, 0.1% NP-40). After washes, 200 µl of reaction buffer (20 mM HEPES pH 7.4, 2 mM MgCl_2_, 0.1 mg/ml bovine serum albumin, mM spermidine, 0.1% NP-40) with 0.5 U/μl RNaseOUT (Invitrogen), 0.5 μg/μl yeast total RNA and 0 to 100 nM of Flag-TOE1 was added to the isolated snRNPs. Reaction tubes were incubated with gentle rotation at 30 °C for 30 minutes followed by incubation at 4°C for 30 minutes. The beads were then isolated using a magnetic stand and resuspended in 100 µl reaction buffer with 100 µl formamide/2 mM EDTA and incubated at 80°C for 5 minutes. 1 ml of TRIzol (Thermo Fisher) was added to each sample. RNA extraction, 3’ end sequencing library preparation, sequencing, and data analysis was performed as described above.

### In vitro transcription and 3’ end processing assay

TMG-, m7G-capped, or uncapped U1 snRNAs were produced by *in vitro* transcription using T7 RNA polymerase (NEB). Briefly, 500 ng of DNA template, produced by PCR from a U1 snRNA expression plasmid^57^ using T7_U1-Forward primer with T7_U1-Reverse_20geno or T7_U1-Reverse-10A primers (Supplementary Table S1), was mixed with 8 mM m7G cap analog (ARCA; NEB), TMG cap analog (Jena Bioscience) or no cap analog, 0.05 M DTT, 1 μl RNaseOUT (2 U/µl; Thermo Fisher), 2 μl NTP buffer mix (1.34 mM final concentration for each NTP, NEB), 2 μl T7 RNA Polymerase Mix (NEB) in a total of 20 μl. *In vitro* transcription was performed at 37°C for 16 hours. RNA products were purified using an RNA Clean & Concentrator kit (Zymo Research) and quantified using a Nanodrop (Thermo Fisher) per manufacturer’s recommendations. For TOE1 3’ end processing assays, 10 ng of RNA was mixed with 100 nM Flag-tagged TOE1 in 10 μl reaction buffer containing 20 mM HEPES pH 7.4, 2 mM MgCl_2_, 100 μg/ml BSA, 1 mM Spermidine, 0.1% NP-40, 0.5 U/μl RNaseOUT, 0.5 μg/μl yeast total RNA. The processing reaction was performed at 37°C for 0 to 120 minutes. Reactions were terminated with 10 µl of denaturing load buffer (95% formamide, 10 mM EDTA, 0.01% Bromophenol Blue, 0.01% Xylene Cyanol) followed by incubation at 80°C for 10 minutes to denature the RNA. The reaction products were subsequently separated in a 7% acrylamide/6M urea denaturing gel. RNA was stained with Sybr Gold (Thermo Fisher) by nutation in the dark for 30 mins. RNA was imaged by a Typhoon gel imager (Amersham). Gel images were analyzed using ImageJ.

### qPCR assays

AR17 (Supplementary Table S1)-primed cDNA was amplified using Fast SYBR Green master mix (Thermo Fisher) with primers for RNAs of interest (Supplementary Table S1) on a QuantStudio Real-Time PCR system (Thermo Fisher). Relative levels were quantified using the ΔΔC_t_ method^73^.

### Western blotting

Western blots were performed by separating proteins in SDS-polyacrylamide gels followed by transfer to nitrocellulose membranes using standard procedures. Membranes were incubated overnight at 4°C with rabbit polyclonal anti-CAF1Z/TOE1^20^ at 1:1,000, rabbit polyclonal anti-UPF1^74^ at 1:1,000, mouse monoclonal anti-SNRPB (Thermo Fisher) at 1:500, each in PBS with 0.1% Tween (PBST) and 5% nonfat milk, or with mouse monoclonal anti-U1-70K (Synaptic System) at 1:1,000 in PBST with 5% BSA. Secondary antibodies were goat anti-rabbit IRDye 680RD (LI-COR) at 1:15,000, HPR donkey anti-rabbit IgG (H+L) (Thermo Fisher), or HPR goat anti-mouse rabbit IgG (H+L) (Thermo Fisher) at 1:20,000 in PBST with 5% nonfat milk. Western blots were visualized using an Odyssey Fc imaging system (LI-COR).

## Supplementary Figure Legends

**Supplementary Figure S1 related to Figure 1.**
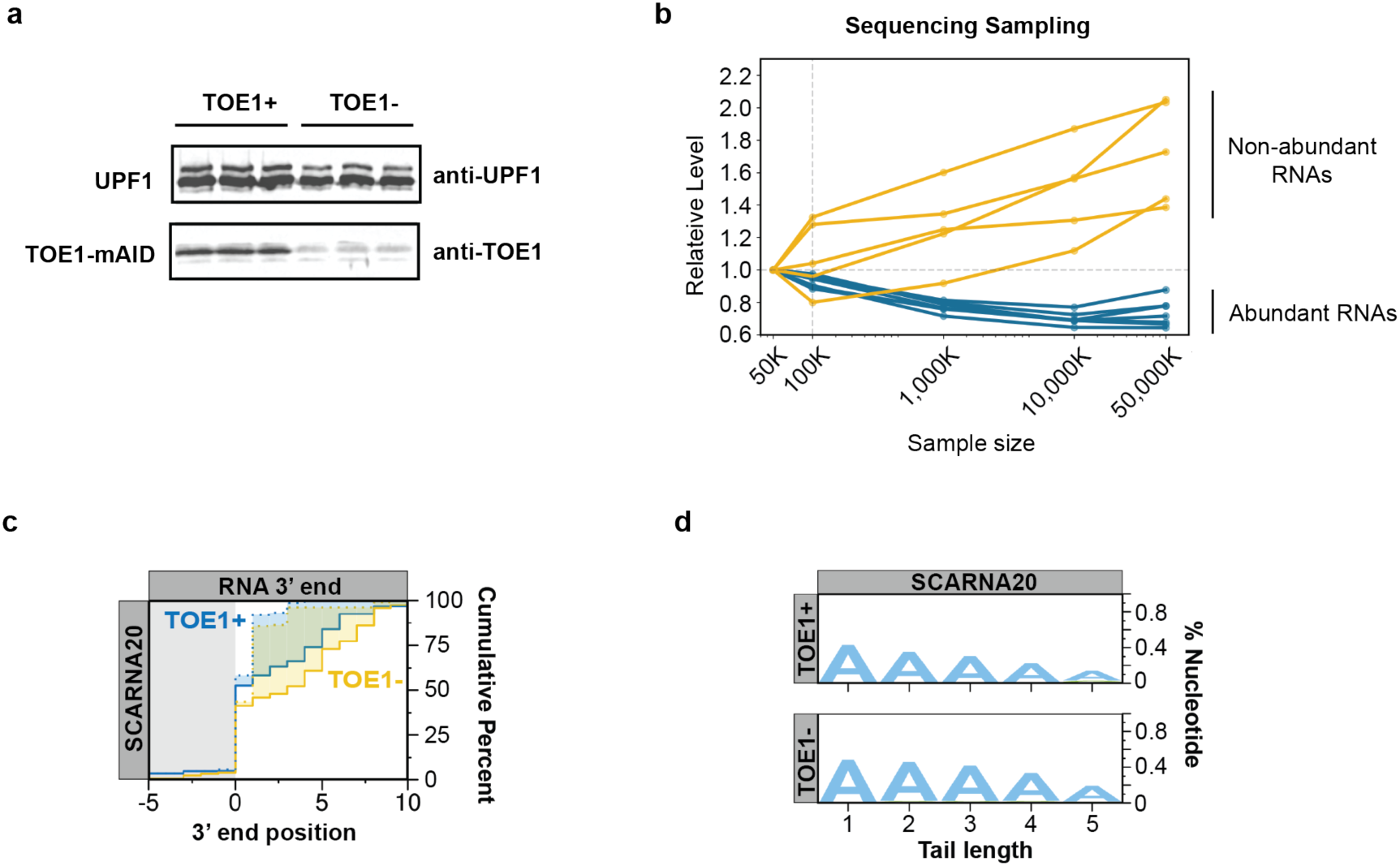
TOE1 shows specificity towards Pol II snRNAs. (**a**) Western blots showing depletion of TOE1. UPF1 serves as a loading control. **(b)** Plot showing the fraction of non-duplicate mapped reads for abundant small ncRNAs (RNU1, RNU2, RNU2-2P, SNORD3, RN7SL1, RN7SL2, RN7SL3, RN7SK) compared with select low abundance RNAs (SNORD13, SNORA71A, SNORA70, SNORA77B, SNORA71C) following sampling of 50k to 50,000k total reads, as normalized to sampling of 50k total reads. The vertical dashed line shows the sampling size (100k reads) used for analyses of the shown abundant small ncRNAs. All other RNAs were analyzed from the full library. **(c)** Cumulative plot showing distributions of 3’ end positions for SCARNA20 in control (TOE1+, blue) and TOE1-depleted (TOE1-, yellow) conditions. **(d)** Sequence logo plots representing the percentage of post-transcriptionally added nucleotides for SCARNA20 under control (TOE1+) or TOE1-depleted (TOE1-) conditions.

**Supplementary Figure S2 related to Figure 2.**
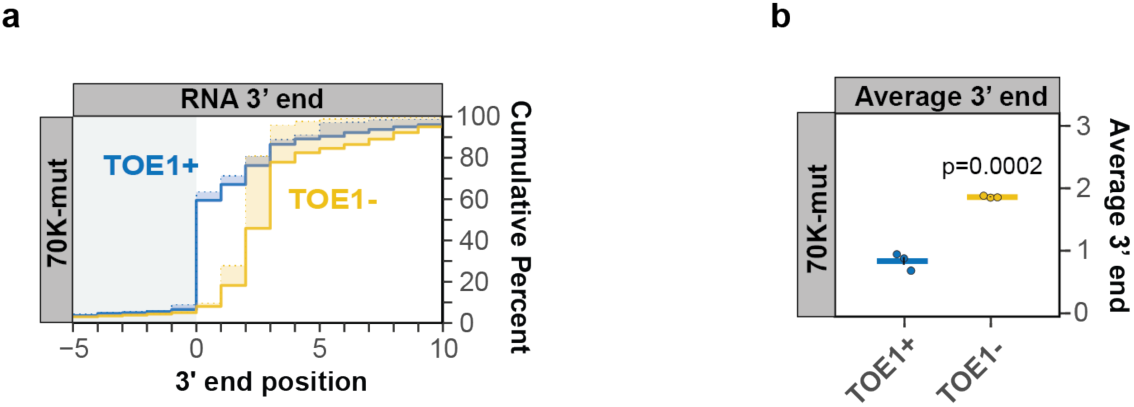
Sm complex– and U1-70K-binding motifs of U1 snRNA promote 3’ end processing. (**a**) Cumulative plot showing 3’ end distributions for the 70K-mutant U1 snRNA under TOE1+ (blue) or TOE1-(yellow) conditions. **(b)** Average 3’ end positions for the 70K mutant U1 snRNA in TOE1+ (blue) and TOE1-(yellow) conditions.

**Supplementary Figure S3 related to Figure 3.**
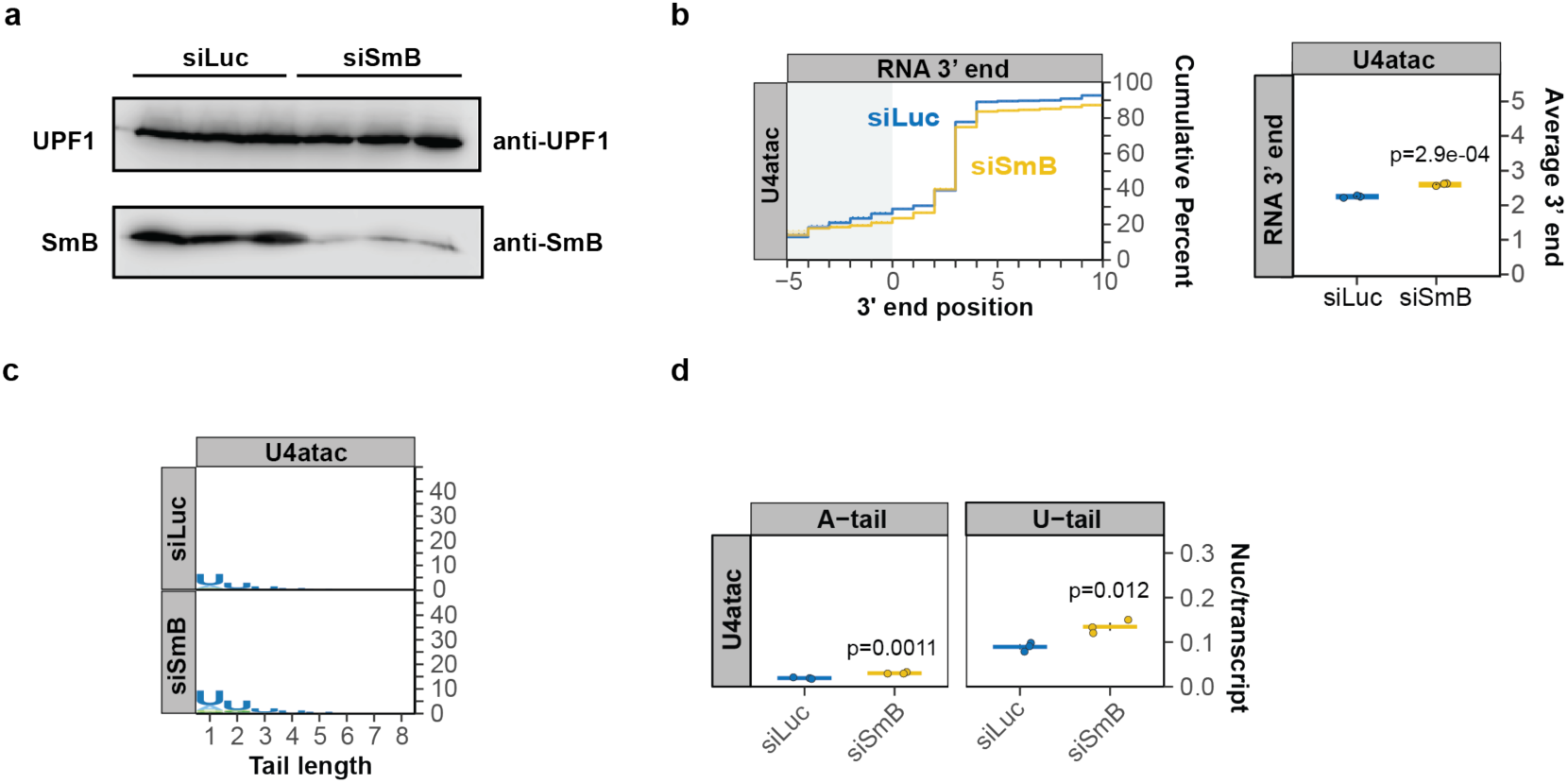
Sm complex depletion impairs 3’ end processing of multiple Pol II snRNAs. (**a**) Western blot showing depletion of SmB. UPF1 serves as a loading control. **(b)** Left, cumulative plots showing 3’ end distributions of nascent minor class Pol II snRNA U4atac in control (siLuc; blue) versus SmB (siSmB; yellow) knock down conditions. The average of three individual biological repeats is plotted. Right, average 3’ end lengths for U4atac snRNA. Dots represent individual biological repeats and SEMs are shown as vertical black lines. P-values were calculated by Student’s two-tailed t-test. **(c)** Sequence logo plots representing the percentage of post-transcriptionally added nucleotides for U4atac snRNA under control (siLuc) or SmB knock down (siSmB) conditions. **(d)** Average number of post-transcriptionally added adenosine or uridine nucleotides per transcript for U4atac snRNA, plotted as in panel *b*.

**Supplementary Figure S4 related to Figure 4.**
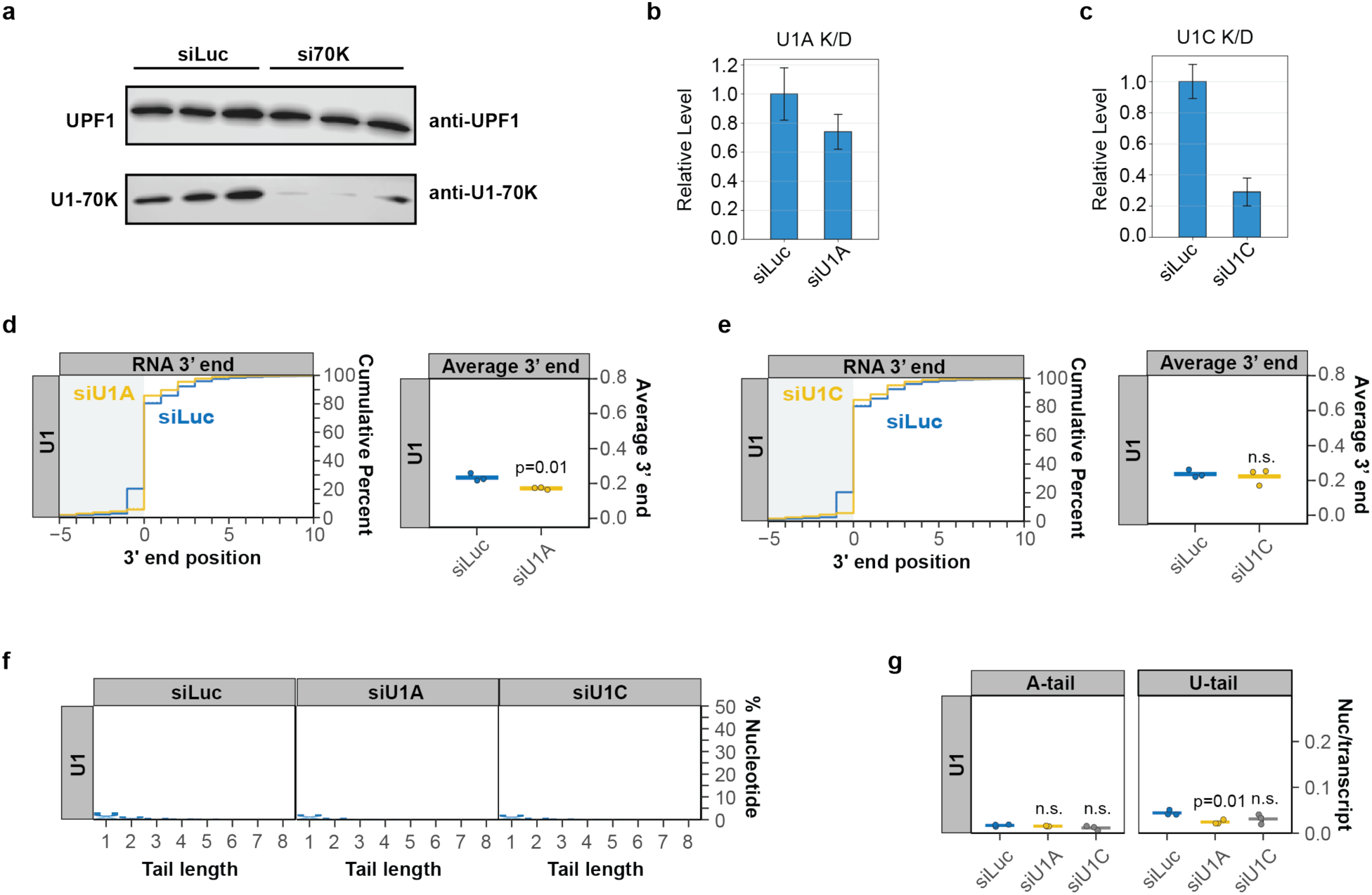
U1-70K stimulates U1 snRNA 3’ end processing by promoting Sm complex assembly. (**a**) Western blot showing depletion of U1-70K. UPF1 serves as a loading control. **(b, c)** Relative levels of U1A (panel *b*) and U1C (panel *c*) mRNAs in U1A– and U1C-depleted conditions, respectively, compared to control (siLuc) conditions, measured by RT-qPCR. Samples were normalized to the average level of GAPDH and Mitochondrial 12S rRNA. Error bars represent SEMs from three individual experiments. **(d, e)** Left, cumulative plots showing 3’ end distributions of nascent U1 snRNA in control (siLuc; blue) versus U1A (panel *d*) or U1C (panel *e*) (yellow) knock-down conditions. Right, average 3’ end lengths for U1 snRNA. Dots represent individual biological repeats and SEMs are shown as vertical black lines. P-values were calculated by Student’s two-tailed t-test. **(f)** Sequence logo plots representing the percentage of post-transcriptionally added nucleotides for nascent U1 snRNA under control (siLuc), U1A knock-down (siU1A), and U1C knock-down (siU1C) conditions. **(g)** Average number of post-transcriptionally added adenosine or uridine nucleotides per transcript, plotted as in panels *d* and *e*.

**Supplementary Figure S5 related to Figure 6.**
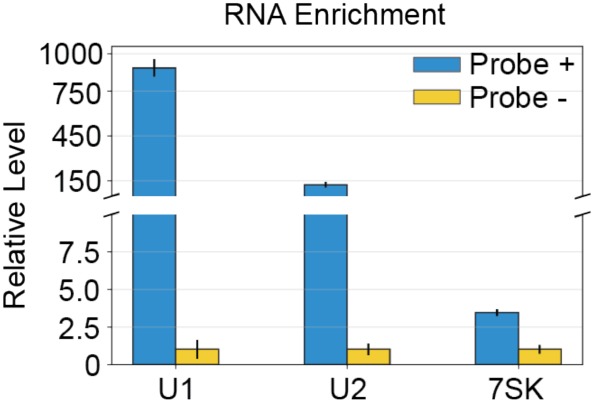
TOE1 directly recognizes the Sm complex-assembled U1 snRNP. Bar plots showing the relative enrichment of U1, U2, and 7SK RNAs following 2’-OMe-RNA oligonucleotide pull-down, measured by RT-qPCR, in pull-down (probe+, Blue) and negative control (probe-, yellow) conditions. Mitochondrial 12S rRNA served as an internal control.

**Supplementary Figure S6 related to Figure 7.**
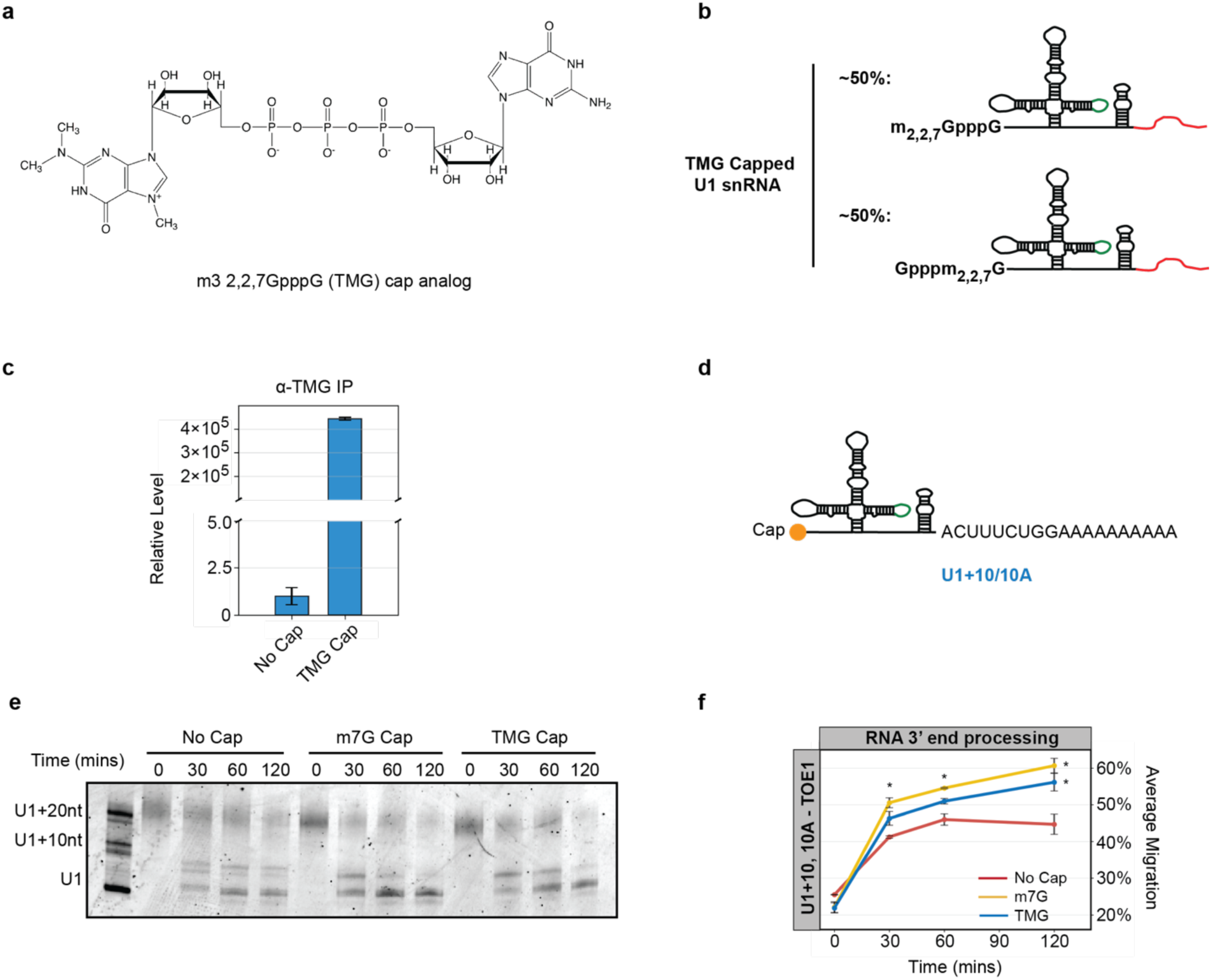
U1 snRNA processing by TOE1 is activated by the 5’ TMG cap. (**a**) Structure of the TMG cap analog. **(b)** The TMG-capped U1 snRNA is a mixture of m_2,2,7_GpppG-capped and Gpppm_2,2,7_G-capped U1 snRNA. **(c)** Relative level of *in vitro* transcribed U1 snRNA with TMG cap or no cap following anti-TMG Immunoprecipitation, measured by RT-qPCR. Samples were normalized to input. **(d)** Schematic of *in vitro* transcribed U1 snRNA with a 10-nucleotide genomic-encoded tail followed by a 10-nucleotides oligo(A) tail (U1+10/10A). **(e, f)** Same as Figure 7 panels *b* and *c*, using U1+10/10A RNA as the substrate for TOE1 processing.

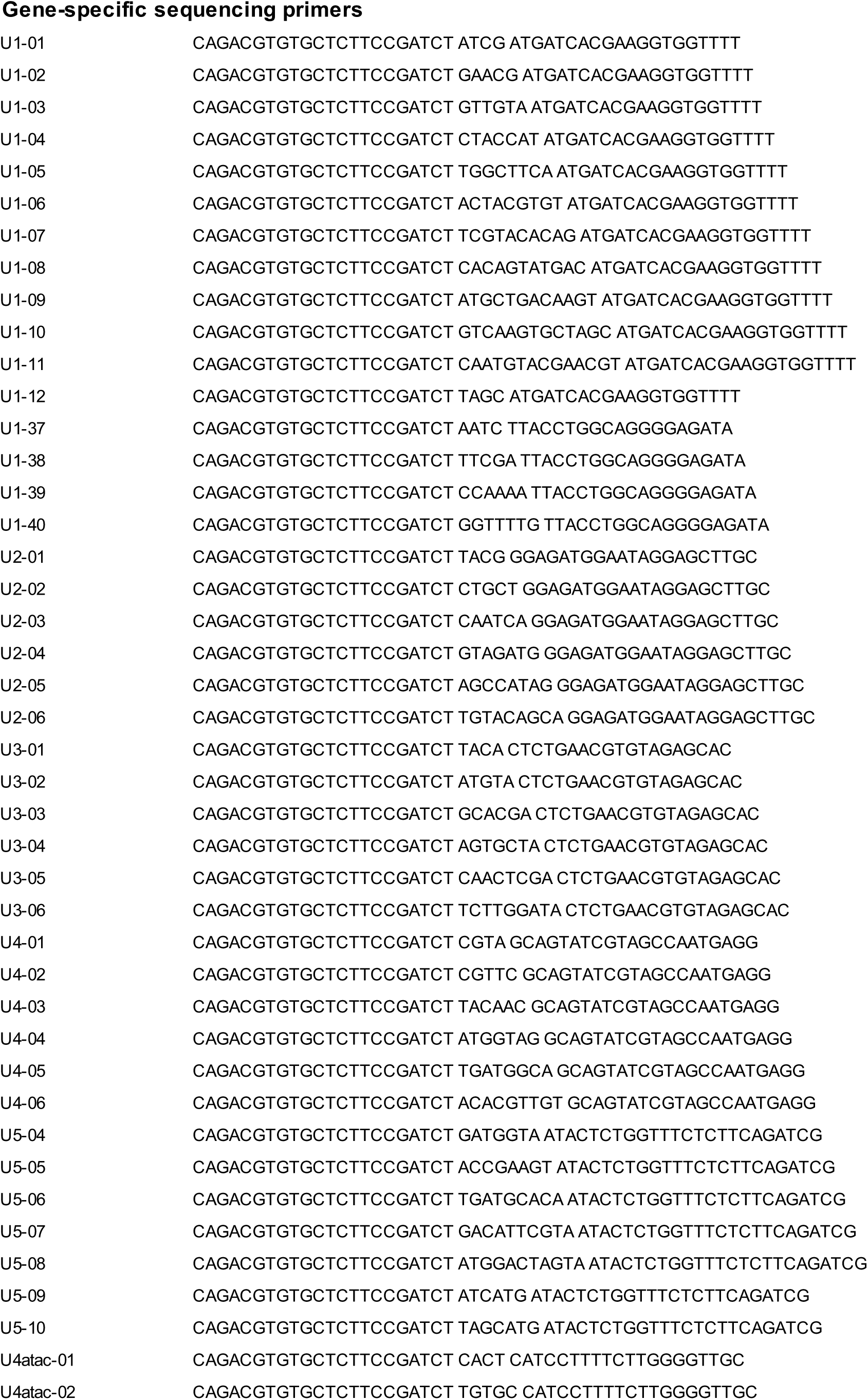
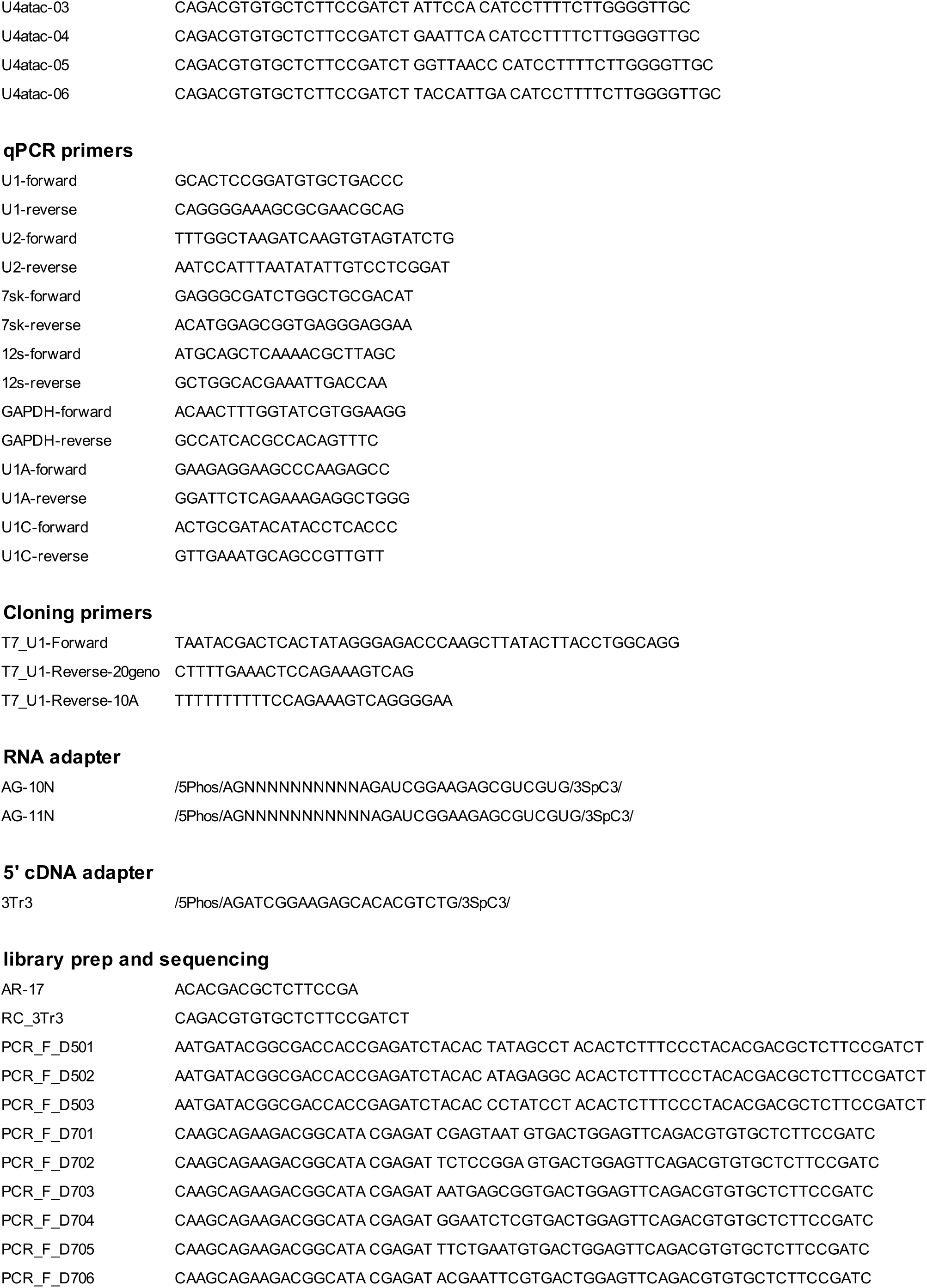
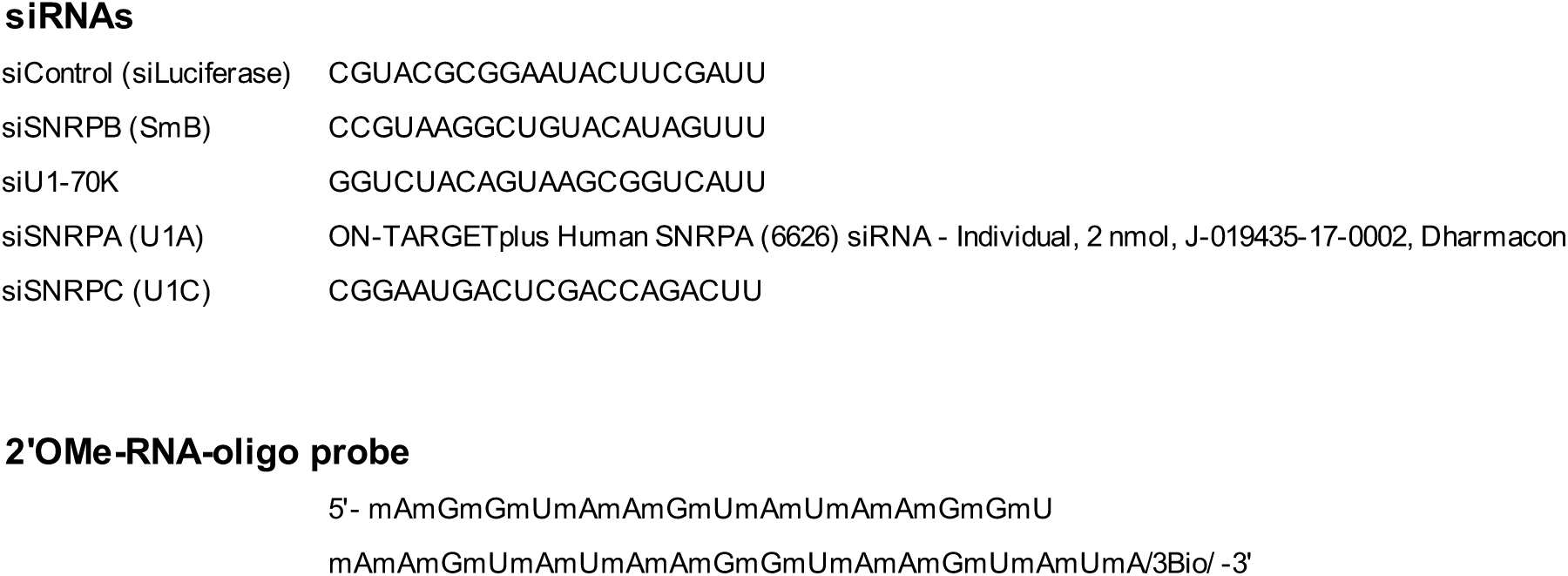
Supplementary TableS1.

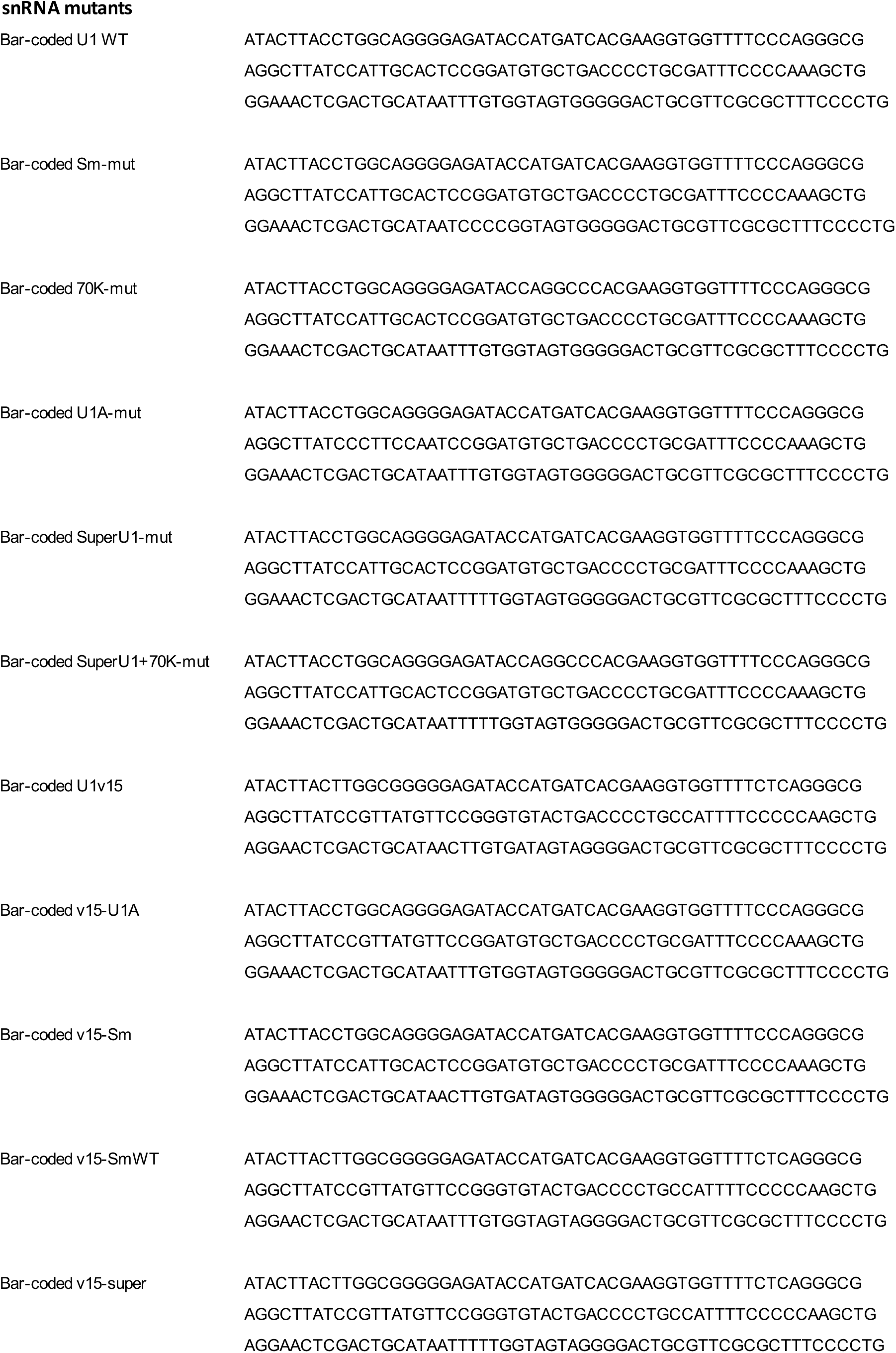
Supplementary TableS2.

## References

1. Christov, C. P., Gardiner, T. J., Szüts, D. & Krude, T. Functional Requirement of Noncoding Y RNAs for Human Chromosomal DNA Replication. Mol. Cell. Biol. 26, 6993–7004 (2006).

2. Nguyen, V. T., Kiss, T., Michels, A. A. & Bensaude, O. 7SK small nuclear RNA binds to and inhibits the activity of CDK9/cyclin T complexes. Nature 414, 322–325 (2001).

3. Yang, Z., Zhu, Q., Luo, K. & Zhou, Q. The 7SK small nuclear RNA inhibits the CDK9/cyclin T1 kinase to control transcription. Nature 414, 317–322 (2001).

4. Zhuang, Y. & Weiner, A. M. A compensatory base change in U1 snRNA suppresses a 5ʹ splice site mutation. Cell 46, 827–835 (1986).

5. Parker, R., Siliciano, P. G. & Guthrie, C. Recognition of the TACTAAC box during mRNA splicing in yeast involves base pairing to the U2-like snRNA. Cell 49, 229–239 (1987).

6. Wightman, B., Ha, I. & Ruvkun, G. Posttranscriptional regulation of the heterochronic gene lin-14 by lin-4 mediates temporal pattern formation in C. elegans. Cell 75, 855–862 (1993).

7. Lafontaine, D. L. J. Noncoding RNAs in eukaryotic ribosome biogenesis and function. Nat. Struct. Mol. Biol. 22, 11–19 (2015).

8. Chanfreau, G., Legrain, P. & Jacquier, A. Yeast RNase III as a key processing enzyme in small nucleolar RNAs metabolism. J. Mol. Biol. 284, 975–988 (1998).

9. Petfalski, E., Dandekar, T., Henry, Y. & Tollervey, D. Processing of the Precursors to Small Nucleolar RNAs and rRNAs Requires Common Components. Mol. Cell. Biol. 18, 1181–1189 (1998).

10. Lardelli, R. M. et al. Biallelic mutations in the 3ʹ exonuclease TOE1 cause pontocerebellar hypoplasia and uncover a role in snRNA processing. Nat. Genet. 49, 457–464 (2017).

11. Son, A., Park, J.-E. & Kim, V. N. PARN and TOE1 Constitute a 3ʹ End Maturation Module for Nuclear Non-coding RNAs. Cell Rep. 23, 888–898 (2018).

12. Lardelli, R. M. & Lykke-Andersen, J. Competition between maturation and degradation drives human snRNA 3ʹ end quality control. Genes Dev. 34, 989–1001 (2020).

13. Fatica, A. Yeast snoRNA accumulation relies on a cleavage-dependent/polyadenylation-independent 3’-processing apparatus. EMBO J. 19, 6218–6229 (2000).

14. Allmang, C. Functions of the exosome in rRNA, snoRNA and snRNA synthesis. EMBO J. 18, 5399–5410 (1999).

15. Van Hoof, A., Lennertz, P. & Parker, R. Yeast Exosome Mutants Accumulate 3ʹ-Extended Polyadenylated Forms of U4 Small Nuclear RNA and Small Nucleolar RNAs. Mol. Cell. Biol. 20, 441–452 (2000).

16. Kawamoto, T., Yoshimoto, R., Taniguchi, I., Kitabatake, M. & Ohno, M. ISG20 and nuclear exosome promote destabilization of nascent transcripts for spliceosomal U snRNAs and U1 variants. Genes Cells 26, 18–30 (2021).

17. Ustianenko, D. et al. TUT-DIS3L2 is a mammalian surveillance pathway for aberrant structured non-coding RNAs. EMBO J. 35, 2179–2191 (2016).

18. Łabno, A. et al. Perlman syndrome nuclease DIS3L2 controls cytoplasmic non-coding RNAs and provides surveillance pathway for maturing snRNAs. Nucleic Acids Res. gkw649 (2016) doi:10.1093/nar/gkw649.

19. Roithová, A., Feketová, Z., Vaňáčová, Š. & Staněk, D. DIS3L2 and LSm proteins are involved in the surveillance of Sm ring-deficient snRNAs. Nucleic Acids Res. 48, 6184–6197 (2020).

20. Wagner, E., Clement, S. L. & Lykke-Andersen, J. An Unconventional Human Ccr4-Caf1 Deadenylase Complex in Nuclear Cajal Bodies. Mol. Cell. Biol. 27, 1686–1695 (2007).

21. Körner, C. G. & Wahle, E. Poly(A) Tail Shortening by a Mammalian Poly(A)-specific 3ʹ-Exoribonuclease. J. Biol. Chem. 272, 10448–10456 (1997).

22. Berndt, H. et al. Maturation of mammalian H/ACA box snoRNAs: PAPD5-dependent adenylation and PARN-dependent trimming. RNA 18, 958–972 (2012).

23. Moon, D. H. et al. Poly(A)-specific ribonuclease (PARN) mediates 3ʹ-end maturation of the telomerase RNA component. Nat. Genet. 47, 1482–1488 (2015).

24. Shukla, S., Schmidt, J. C., Goldfarb, K. C., Cech, T. R. & Parker, R. Inhibition of telomerase RNA decay rescues telomerase deficiency caused by dyskerin or PARN defects. Nat. Struct. Mol. Biol. 23, 286–292 (2016).

25. Roake, C. M. et al. Disruption of Telomerase RNA Maturation Kinetics Precipitates Disease. Mol. Cell 74, 688–700.e3 (2019).

26. Deng, T. et al. TOE1 acts as a 3ʹ exonuclease for telomerase RNA and regulates telomere maintenance. Nucleic Acids Res. 47, 391–405 (2019).

27. Izumi, N. et al. Identification and Functional Analysis of the Pre-piRNA 3ʹ Trimmer in Silkworms. Cell 164, 962–973 (2016).

28. Anastasakis, D. et al. Mammalian PNLDC1 is a novel poly(A) specific exonuclease with discrete expression during early development. Nucleic Acids Res. 44, 8908–8920 (2016).

29. Ding, D. et al. PNLDC1 is essential for piRNA 3ʹ end trimming and transposon silencing during spermatogenesis in mice. Nat. Commun. 8, 819 (2017).

30. Mroczek, S., et al. *C16orf57*, a gene mutated in poikiloderma with neutropenia, encodes a putative phosphodiesterase responsible for the U6 snRNA 3ʹ end modification. Genes Dev. 26, 1911–1925 (2012).

31. Shchepachev, V., Wischnewski, H., Missiaglia, E., Soneson, C. & Azzalin, C. M. Mpn1, Mutated in Poikiloderma with Neutropenia Protein 1, Is a Conserved 3ʹ-to-5ʹ RNA Exonuclease Processing U6 Small Nuclear RNA. Cell Rep. 2, 855–865 (2012).

32. Jeong, H.-C., et al. USB1 is a miRNA deadenylase that regulates hematopoietic development. (2023).

33. Tummala, H. et al. Poly(A)-specific ribonuclease deficiency impacts telomere biology and causes dyskeratosis congenita. J. Clin. Invest. 125, 2151–2160 (2015).

34. Dhanraj, S. et al. Bone marrow failure and developmental delay caused by mutations in poly(A)-specific ribonuclease (*PARN*). J. Med. Genet. 52, 738–748 (2015).

35. Nagirnaja, L. et al. Variant *PNLDC1*, Defective piRNA Processing, and Azoospermia. N. Engl. J. Med. 385, 707–719 (2021).

36. Walne, A. J., Vulliamy, T., Beswick, R., Kirwan, M. & Dokal, I. Mutations in C16orf57 and normal-length telomeres unify a subset of patients with dyskeratosis congenita, poikiloderma with neutropenia and Rothmund–Thomson syndrome. Hum. Mol. Genet. 19, 4453–4461 (2010).

37. Will, C. L. & Luhrmann, R. Spliceosome Structure and Function. Cold Spring Harb. Perspect. Biol. 3, a003707–a003707 (2011).

38. Baillat, D. et al. Integrator, a Multiprotein Mediator of Small Nuclear RNA Processing, Associates with the C-Terminal Repeat of RNA Polymerase II. Cell 123, 265–276 (2005).

39. Salditt-Georgieff, M., Harpold, M., Chen-Kiang, S. & Darnell, J. E. The addition of 5ʹ cap structures occurs early in hnRNA synthesis and prematurely terminated molecules are capped. Cell 19, 69–78 (1980).

40. Ohno, M., Segref, A., Bachi, A., Wilm, M. & Mattaj, I. W. PHAX, a Mediator of U snRNA Nuclear Export Whose Activity Is Regulated by Phosphorylation. Cell 101, 187–198 (2000).

41. So, B. R. et al. A U1 snRNP–specific assembly pathway reveals the SMN complex as a versatile hub for RNP exchange. Nat. Struct. Mol. Biol. 23, 225–230 (2016).

42. Fischer, U., Liu, Q. & Dreyfuss, G. The SMN–SIP1 Complex Has an Essential Role in Spliceosomal snRNP Biogenesis. Cell 90, 1023–1029 (1997).

43. Liu, Q., Fischer, U., Wang, F. & Dreyfuss, G. The Spinal Muscular Atrophy Disease Gene Product, SMN, and Its Associated Protein SIP1 Are in a Complex with Spliceosomal snRNP Proteins. Cell 90, 1013–1021 (1997).

44. Buhler, D., Raker, V., Luhrmann, R. & Fischer, U. Essential Role for the Tudor Domain of SMN in Spliceosomal U snRNP Assembly: Implications for Spinal Muscular Atrophy. Hum. Mol. Genet. 8, 2351–2357 (1999).

45. Friesen, W. J. et al. The Methylosome, a 20S Complex Containing JBP1 and pICln, Produces Dimethylarginine-Modified Sm Proteins. Mol. Cell. Biol. 21, 8289–8300 (2001).

46. Meister, G., Bühler, D., Pillai, R., Lottspeich, F. & Fischer, U. A multiprotein complex mediates the ATP-dependent assembly of spliceosomal U snRNPs. Nat. Cell Biol. 3, 945–949 (2001).

47. Pellizzoni, L., Yong, J. & Dreyfuss, G. Essential Role for the SMN Complex in the Specificity of snRNP Assembly. Science 298, 1775–1779 (2002).

48. Massenet, S., Pellizzoni, L., Paushkin, S., Mattaj, I. W. & Dreyfuss, G. The SMN Complex Is Associated with snRNPs throughout Their Cytoplasmic Assembly Pathway. Mol. Cell. Biol. 22, 6533–6541 (2002).

49. Yong, J., Kasim, M., Bachorik, J. L., Wan, L. & Dreyfuss, G. Gemin5 Delivers snRNA Precursors to the SMN Complex for snRNP Biogenesis. Mol. Cell 38, 551–562 (2010).

50. Mattaj, I. W. Cap trimethylation of U snRNA is cytoplasmic and dependent on U snRNP protein binding. Cell 46, 905–911 (1986).

51. Mouaikel, J., Verheggen, C., Bertrand, E., Tazi, J. & Bordonné, R. Hypermethylation of the Cap Structure of Both Yeast snRNAs and snoRNAs Requires a Conserved Methyltransferase that Is Localized to the Nucleolus. Mol. Cell 9, 891–901 (2002).

52. Mouaikel, J. et al. Interaction between the small-nuclear-RNA cap hypermethylase and the spinal muscular atrophy protein, survival of motor neuron. EMBO Rep. 4, 616–622 (2003).

53. Huber, J. et al. Snurportin1, an m3G-cap-specific nuclear import receptor with a novel domain structure. EMBO J. 17, 4114–4126 (1998).

54. Palacios, I. Nuclear import of U snRNPs requires importin beta. EMBO J. 16, 6783–6792 (1997).

55. Bohnsack, M. T. & Sloan, K. E. Modifications in small nuclear RNAs and their roles in spliceosome assembly and function. Biol. Chem. 399, 1265–1276 (2018).

56. Yang, H., Moss, M. L., Lund, E. & Dahlberg, J. E. Nuclear Processing of the 3ʹ-Terminal Nucleotides of Pre-U1 RNA in *Xenopus laevis* Oocytes. Mol. Cell. Biol. 12, 1553–1560 (1992).

57. Shukla, S. & Parker, R. Quality control of assembly-defective U1 snRNAs by decapping and 5ʹ-to-3ʹ exonucleolytic digestion. Proc. Natl. Acad. Sci. 111, (2014).

58. Tsai, D. E., Harper, D. S. & Keene, J. D. U1-snRNP-A protein selects a ten nucleotide consensus sequence from a degenerate RNA pool presented in various structural contexts. Nucleic Acids Res. 19, 4931–4936 (1991).

59. Hernández, H. et al. Isoforms of U1-70k Control Subunit Dynamics in the Human Spliceosomal U1 snRNP. PLoS ONE 4, e7202 (2009).

60. Martıńez, J., et al. A 54-kDa Fragment of the Poly(A)-specific Ribonuclease Is an Oligomeric, Processive, and Cap-interacting Poly(A)-specific 3ʹ Exonuclease. J. Biol. Chem. 275, 24222–24230 (2000).

61. Gao, M., Fritz, D. T., Ford, L. P. & Wilusz, J. Interaction between a Poly(A)-Specific Ribonuclease and the 5ʹ Cap Influences mRNA Deadenylation Rates In Vitro. Mol. Cell 5, 479–488 (2000).

62. Dehlin, E., Wormington, M., Körner, C. G. & Wahle, E. Cap-dependent deadenylation of mRNA. EMBO J. 19, 1079–1086 (2000).

63. Cheng, L. et al. Loss of the RNA trimethylguanosine cap is compatible with nuclear accumulation of spliceosomal snRNAs but not pre-mRNA splicing or snRNA processing during animal development. PLOS Genet. 16, e1009098 (2020).

64. Chen, L. et al. TGS1 impacts snRNA 3ʹ-end processing, ameliorates *survival motor neuron* – dependent neurological phenotypes *in vivo* and prevents neurodegeneration. Nucleic Acids Res. 50, 12400–12424 (2022).

65. O’Reilly, D. et al. Differentially expressed, variant U1 snRNAs regulate gene expression in human cells. Genome Res. 23, 281–291 (2013).

66. Mabin, J. W., Lewis, P. W., Brow, D. A. & Dvinge, H. Human spliceosomal snRNA sequence variants generate variant spliceosomes. RNA 27, 1186–1203 (2021).

67. Chen, L. et al. Loss of Human TGS1 Hypermethylase Promotes Increased Telomerase RNA and Telomere Elongation. Cell Rep. 30, 1358–1372.e5 (2020).

68. Kufel, J. & Grzechnik, P. Small Nucleolar RNAs Tell a Different Tale. Trends Genet. 35, 104– 117 (2019).

69. Dobin, A. et al. STAR: ultrafast universal RNA-seq aligner. Bioinformatics 29, 15–21 (2013).

70. Quinlan, A. R. & Hall, I. M. BEDTools: a flexible suite of utilities for comparing genomic features. Bioinformatics 26, 841–842 (2010).

71. Li, H. et al. The Sequence Alignment/Map format and SAMtools. Bioinformatics 25, 2078– 2079 (2009).

72. Nicholson-Shaw, T. & Lykke-Andersen, J. Tailer: A Pipeline for Sequencing-Based Analysis of Non-Polyadenylated RNA 3’ End Processing.

73. Livak, K. J. & Schmittgen, T. D. Analysis of Relative Gene Expression Data Using Real-Time Quantitative PCR and the 2−ΔΔCT Method. Methods 25, 402–408 (2001).

74. Lykke-Andersen, J., Shu, M.-D. & Steitz, J. A. Human Upf Proteins Target an mRNA for Nonsense-Mediated Decay When Bound Downstream of a Termination Codon. Cell 103, 1121–1131 (2000).

